# Sustained Isobutene Production by *Synechocystis* sp. PCC 6803 Entrapped in Polyvinyl Alcohol Hydrogel Beads

**DOI:** 10.1101/2025.01.02.631087

**Authors:** Sindhujaa Vajravel, Sanjukta Aravind, Karin Stensjö

## Abstract

Cyanobacteria convert CO_2_ into valuable compounds using solar energy, making them ideal for sustainable isobutene production, a key precursor for fuels and chemicals. This study aimed to enhance isobutene production in engineered *Synechocystis* sp. PCC 6803 strains: Syn- *Rn*KICD, producing isobutene from α-ketoisocaproate (KIC) via *Rattus norvegicus* α- ketoisocaproate dioxygenase (*Rn*KICD), and Syn-F336V, expressing a mutant variant of *Rn*KICD with improved KIC specificity. We investigated the effects of varying culture conditions, including light intensity, inorganic carbon, and nitrogen on isobutene production. Nitrogen limitation emerged as a critical factor, improving yields by reducing growth. This likely occurred due to decreased competition from branched-chain amino acid (BCAA) biosynthesis, redirecting carbon toward isobutene synthesis. However, prolonged nitrogen limitation ultimately reduced productivity due to impaired metabolic functions. To address this limitation, we employed a polyvinyl alcohol-sodium alginate (PVA-SA) hydrogel, crosslinked with B(OH)₄⁻ and Ca²⁺, to entrap cells. This approach restricted growth while maintaining cell viability and isobutene productivity. Optimizing crosslinking parameters such as time, pH, and the hydrogel-to-cell mass ratio improved bead stability under bicarbonate and nitrate supply. This strategy extended cell viability and isobutene productivity in Syn-*Rn*KICD and Syn- F336V by nearly a month, increasing yields by 60% and 80%, respectively, compared to suspension cells, achieving a maximum yield of 94 mg/g DW. This study underscores the importance of optimizing environmental conditions for isobutene production in *Synechocystis* and highlights the effectiveness of PVA-SA cell entrapment as a biocatalyst platform for sustained chemical production.

**Highlights:**

- Optimized light, carbon, and nitrogen sources enhanced isobutene production.
- Rate of isobutene production increased 10-fold compared to our previous report.
- Nitrogen limitation improved isobutene yield by redirecting carbon from growth.
- PVA-entrapped cells sustained isobutene productivity for nearly 31 days.

## 1. Introduction

As global efforts to reduce reliance on fossil resources intensify, the push for renewable chemical production has become crucial for achieving sustainability and protecting the environment. Photosynthetic cyanobacteria are being recognized for their potential in sustainable chemical production, efficiently using sunlight and converting atmospheric CO₂ to synthesize valuable organic compounds. These microorganisms exhibit notable solar energy conversion efficiencies, reaching ∼8–10% for solar-to-biomass conversion, surpassing the typical 4-6% efficiency observed in terrestrial plants (Melis, 2009; Zhu et al., 2008). Additionally, their rapid growth rates, high lipid content for biofuel production, and ability to thrive in diverse environments, including polluted and wastewater for potential bioremediation applications, make cyanobacteria efficient and environmentally friendly resources (Farrokh et al., 2019).

Cyanobacteria’s genetic flexibility offers extensive bioengineering potential to optimize photosynthetic pathways for efficient conversion of CO₂ into targeted organic molecules. Advances in synthetic biology and metabolic engineering have greatly expanded their capacity to produce a diverse array of chemicals. These include fuels such as ethanol and butanol, petrochemical industrial precursors like isoprene, isobutene, and ethylene, and other high-value compounds such as 2,3-butanediol, sucrose, lactic acid, isobutyraldehyde, and bisabolene (Angermayr et al., 2015; Carroll et al., 2018; Santos-Merino et al., 2019). Furthermore, isoprene produced by photosynthetic cyanobacteria can be converted into kerosene-type fuel precursors through photochemical dimerization processes, highlighting the significant potential of these organisms for producing renewable, high-energy-density fuels (Rana et al., 2022). Additionally, the ability of these small, volatile hydrocarbons to easily diffuse out of cyanobacterial cell membranes without damaging the cells enhances the economic and energy efficiency of the product harvesting process (Lindberg et al., 2010; Ducat et al., 2011).

Isobutene (C₄H₈) is a highly versatile hydrocarbon pivotal in the chemical industry, primarily due to its reactive double bond, which facilitates a variety of photochemical reactions, including oligomerization and polymerization, resulting in a wide range of industrially important products. Its oligomerization potential extends to creating high-density fuels, showcasing its viability in sustainable aviation fuel solutions (Hauge et al., 2005; Taylor et al., 2010; Peters and Taylor, 2013; Kostjuk 2015; Nicholas, 2017). Remarkably, isobutene can undergo visible light induced cationic photopolymerization, producing significant polymers such as plastics and rubbers (Hulnik et al., 2023), underscoring its role in advanced materials science. These chemical reactions are particularly advantageous because they occur under mild conditions, offering significant benefits over traditional thermal processes. Despite these advancements, isobutene is still primarily sourced from fossil fuels, highlighting a significant opportunity and need for developing renewable isobutene production methods to meet the demands of the petrochemical industry sustainably.

Global demand for sustainable isobutene is surging, with the market projected to surpass USD 30.71 billion and 16 million tons by 2024 (Rebolledo-Leiva et al., 2022), reaching USD 38.4 billion by 2030 (Grand View Research). The highest bioisobutene productivity reported so far is 0.45 mg L^-1^ h^-1^ by the yeast *Rhodotorula minut*a (Wilson et al., 2018), but to achieve economic viability, yields of ∼2–4 g L^-1^ h^-1^ are necessary (van Leeuwen et al., 2012). Recent advancements focus on exploiting metabolically modified photosynthetic cyanobacteria to meet these economic and sustainability targets. Our team has engineered the cyanobacterium *Synechocystis* sp. PCC 6803 (hereafter *Synechocystis)* to produce isobutene directly from CO₂ by heterologously expressing the *Rn*KICD, re-routing the central metabolism from pyruvate through the L-leucine biosynthesis pathway-derived metabolite KIC to isobutene, achieving a rate of 91 ng l⁻¹ OD_750_⁻¹ h⁻¹ (Mustila et al., 2021). The mammalian enzyme KICD, catalyzes the decarboxylation and oxygenation of KIC to form β-hydroxy-β- methyl butyrate (HMB) in the cytosol of liver cells as part of the L-leucine catabolic pathway (Sabourin & Bieber 1982). Thus, the *Rn*KICD acts on multiple substrates, catalyzing the conversion of 4-hydroxyphenylpyruvate (HPP) to homogentisate (HGA) and KIC to either HMB or isobutene. Building on these findings, our team recently refined *Rn*KICD’s substrate preference for KIC to further boost isobutene production. Through protein engineering and targeted mutagenesis, a specific mutation, F336V, was identified, which shifts *Rn*KICD*’*s substrate preference toward KIC rather than HPP, significantly enhancing isobutene production (Schumann et al., 2024). However, further optimizations of favorable environmental culture conditions are essential to achieve production levels that meet commercial demands.

The efficiency of photoproduction of high-value compounds by microalgae is significantly influenced by various environmental conditions, including light intensity, pH, temperature, and the availability of macronutrients like inorganic carbon, nitrogen, phosphorus, and sulfur (Miller and Colman, 1980; Gordillo et al., 1998; Aboim et al., 2019; Rodrigues et al., 2023). However, traditional suspension cultures face yield limitations due to excessive biomass accumulation, which increases nutrient demand, oxidative stress, and self-shading effects, ultimately reducing photosynthetic efficiency (Chisti, 2007; Kim et al., 2011; Pedruzi et al., 2019). One promising strategy to address these limitations is the physical entrapment of microalgal cells within hydrogel polymers. One widely used method involves crosslinking sodium alginate (SA) with CaCl₂ to form stable hydrogel beads through ionic bonds between Ca²⁺ ions and the carboxylate groups of guluronic and mannuronic acid residues in alginate chains, resulting in a robust three-dimensional network (Simpson et al., 2004). This whole-cell entrapment approach improved cyanobacterial viability and productivity, as demonstrated by their notable cell viability and enhanced production of ammonia, hydrogen, ethylene, β- phellandrene, and sucrose (Brouers and Hall, 1986; Kosourov et al., 2014; Vajravel et al., 2020; Valsami et al., 2021; Tóth et al., 2022). These improvements were achieved by effective biomass growth mangement, nutrient distribution, enhnced cellular communication, and reduced oxidative stress. However, the use of Ca²⁺ alginate beads in cyanobacterial photoproduction systems presents challenges, particularly due to the potential formation of CaCO₃, which diminishes bead stability, especially in environments where NaHCO₃ is used as a crucial inorganic carbon source in airtight vessels for small volatile compounds production (Rissanen et al., 2022). Therefore, it is important to select suitable hydrogel polymers for the cell entrapment approach that can withstand HCO₃⁻-supplemented cultures.

Polyvinyl alcohol (PVA) hydrogels are widely used in biomedical applications due to their flexible crosslinking capabilities, biocompatibility, biodegradability, and cost-effective (Zhong et al., 2024). PVA can be crosslinked with sodium borate, boric acid, glutaraldehyde, formaldehyde, UV treatment and freeze-thawing (Wu & Wisecarver, 1992; Gohil et al., 2006; Bolto et al., 2009; Hua et al., 2010). When B(OH)₄⁻ ions, formed from H₃BO₃, interact with PVA, they create reversible covalent bonds with the hydroxyl groups of PVA, forming a stable yet dynamic hydrogel structure that can break and reform bonds in response to external forces or environmental changes (Cheng and Rodriguez, 1981; Wu & Wisecarver et al., 1992; Choe et al., 2024; Wu et al., 2024). This dynamic behavior enhances the material’s resilience and adaptability, enabling repeated deformation and recovery, which improves the hydrogel’s functionality in applications such as drug delivery and tissue engineering.

In this study, we aimed to enhance isobutene production in *Synechocystis* strains, Syn- *Rn*KICD and Syn-F336V, by systematically exploring various environmental factors, including light intensity, bicarbonate and nitrogen source availability, and employing borate- PVA hydrogel for cell entrapment. Our findings indicated that incorporating a stabilizing agent such as SA, adjusting the crosslinking duration, maintaining neutral pH, and optimizing the cell biomass-to-hydrogel ratio significantly improved the hydrogel beads’ stability and longevity, where HCO₃ served as a crucial inorganic carbon source. By utilizing this optimized system, we extended the productive phase of isobutene to one month, achieving maximum yields of ∼58 mg/g DW and 94 mg/g DW in the entrapped Syn-*Rn*KICD and Syn-F336V cells, respectively.

## 2. Materials and Methods

### 2.1. Strains and Growth Conditions

The two *Synechocystis* strains were employed in this study were previously constructed: Syn- *Rn*KICD, which produces a KICD enzyme from *Rattus norvegicus* (*Rn*KICD) (Mustila et al., 2021), and Syn-F336V, in which a modified version of *Rn*KICD is generated (Schumann et al., 2024). *Synechocystis* precultures were cultivated photoautotrophically in BG11 liquid medium in 6-well polystyrene plates, illuminated with continuous white light at a photosynthetic photon flux density (PPFD) of 30 µmol photons m⁻² s⁻¹. For the cultivation of experimental cultures, 250 mL Erlenmeyer flasks (E-flasks) containing 80 mL of medium were buffered with 25 mM HEPES-NaOH (pH 7.4) and supplemented with 50 µg/mL kanamycin (Km). The cultures were supplied with 50 mM NaHCO₃ and maintained at 30°C with shaking at 100 rpm under illumination of 50 µmol photons m⁻² s⁻¹. All precultures were initiated from cryopreserved cells stored at -80°C in 10% glycerol.

### 2.2. Isobutene Production and Quantification

For isobutene production, *Synechocystis* cells cultivated in E-flasks were harvested by centrifugation at 5000 rpm for 5 min upon reaching an optical density (OD_750_) of ∼ 1. Following centrifugation, the cells were washed with fresh BG11 medium, adjusted to similar OD_750_ for all cultures, and supplemented with 50 mM NaHCO₃ and 50 µg mL^-1^ Km. A 10 mL aliquot of the suspension cultures was transferred into 20 mL gas-tight vials, which were sealed with Teflon-coated septa.

The suspension cultures were shaken at 100 rpm under an illumination of 50 µmol photons m⁻² s⁻¹ (unless stated otherwise). Every 24 h of post light illumination, the headspace of the vials was sampled through the Teflon septum, and after each injection, the septa were replaced to minimize any loss of produced isobutene. A 5-min interval was allowed during septa replacement to facilitate gas exchange.

Isobutene detection and quantification were performed using an Agilent 7890A Gas Chromatography (GC) system coupled with a 5975 Mass Selective Detector (MSD), equipped with an HP-1ms Ultra I column. Samples were taken from 20 mL GC vials with Teflon-coated septa (VWR) and injected using a PAL-RSI85 autosampler with the PAL-HS2500 gas syringe tool. The GC-MS method for isobutene detection was adapted from Rossoni et al (2015). Prior to injection, GC vials were incubated for 5 min at 40°C to equilibrate the isobutene concentration in the headspace. For detection, 100 μL of each sample were injected at an inlet liner temperature of 150°C with a 100:1 split ratio. The oven temperature was held at 40°C for 3 min before increasing by 30°C/min for 2 min. The retention time for isobutene was ∼1.94 min with an authentic standard (99% v/v C₄H₈, AGA, Sweden). The MS detector operated in selected ion monitoring (SIM) mode to enhance sensitivity, specifically detecting isobutene signals at m/z 41 and 56. Signals within the chromatogram were analyzed for peak area through autointegration. The quantity of isobutene was determined from a calibration curve established using various dilution percentages of a commercial gas standard (99% v/v C₄H₈, AGA, Sweden).

Isobutene productivity was normalized to the initial OD values for suspension cultures and initial cell dry weight (DW) was used for entrapped cells. For a direct comparison of isobutene productivity between suspension and entrapped cells, the initial dry weight (DW) of cells was set to be similar across all samples. The cumulative productivity of isobutene was calculated by summing the isobutene produced at 24-h intervals. The data were averaged from 3 independent measurements taken from 3 replicates (±SD).

### 2.3. Induction of Nitrogen Starvation

Nitrogen (N) starvation in *Synechocystis* cells was induced by omitting NaNO₃ from the BG- 11 medium, the sole N source for the non-nitrogen-fixing cyanobacteria in the E-flask allowing for adaptation as mentioned in the section (2.1). In the experimental cultures, varying N starvation periods were tested to assess the effects of both short-term (24 h) and long-term (72 h) N starvation on *Synechocystis*. After the starvation period the cells were washed with N-free BG-11 medium by centrifuging at 5000 rpm for 5 min (2x). The cells were then prepared for isobutene production by resuspending them in N-free medium and transferring them into tightly sealed vials. They were subsequently adapted under the conditions described in Section 2.2.

### 2.4. Cell entrapment procedures

Syn-*Rn*KICD and Syn-F336V cells were cultivated under experimental conditions as described in Section 2.2. Once the cells reached OD_750_ of 1, the cells were harvested by centrifugation at 5000 rpm for 5 min. Subsequently, the cyanobacterial cells were entrapped in hydrogel for entrapment, as described by Brouers and Hall (1986), with some modifications. We employed a PVA hydrogel crosslinked with H₃BO₃ using a droplet-based dispersion technique via a 2 mL syringe. To form a distinct hydrogel beads, 0.5% SA was incorporated into the 6% PVA hydrogel and crosslinking was achieved using 2% CaCl₂ and 4% H₃BO₃ solutions. We used varying formulations of wet cell biomass and PVA-SA hydrogel, with ratios ranging from 5:6:0.5 to 5:36:3 (cell:PVA:SA) for cell entrapment.

Hydrogel droplets were manually dispensed into the crosslinking agents using a 2 mL syringe. After dispensation, the plates were sealed with Parafilm, and the crosslinking duration was optimized between 5 to 20 h. The instantly formed beads were incubated at 30°C to produce stable beads in the presence of crosslinking agents, under illumination at 30 µmol photons m⁻² s⁻¹. After the desired duration, the beads were thoroughly rinsed with sterile deionized water to remove excess crosslinkers and minimize ionic stress on the cells. The entrapped cells in hydrogel beads were then transferred into 20 mL gas-tight vials containing BG11 medium (with and without NaNO_3_, as stated) with 50 mM NaHCO₃ and subjected to the isobutene production conditions described in Section 2.2.

### 2.5. Measurement of cell density and pigment content

The growth of the cell cultures was assessed spectrophotometrically (Varian 50 Bio UV-Visible Spectrophotometer) by measuring the OD_750_. The chlorophyll *a* (Chl) content was quantified by measuring the absorbance at 665 nm following extraction with 90% methanol (Lichtenthaler, 1987). Additionally, the absorption spectra of the whole cell suspensions were recorded. Prior to measurement, all cultures were standardized to a uniform cell density of OD_750_ ∼ 0.5.

## 3. Results and Discussion

### 3.1. Optimizing isobutene production in Syn-*Rn*KICD: The interplay of light intensity and inorganic carbon

The main strain used in this study; Syn-*Rn*KICD, expresses the *Rn*KICD gene in *Synechocystis* that encodes KICD, an enzyme that has shown to convert KIC to isobutene *in vitro* and *in vivo* (Mustila et al., 2021). To study how varying light intensities affect isobutene production, we conducted experiments under three different photon flux densities: 35 (low light, LL), 85 (moderate light, ML), and 135 (high light, HL) μmol photons m⁻² s⁻¹. To ensure adequate inorganic carbon (Ci) availability for photosynthesis in the air-tight vessels, all cultures were supplemented with 50 mM NaHCO₃. The optimal light intensity for maximizing isobutene production was determined (Fig 1A).

**Figure 1:**
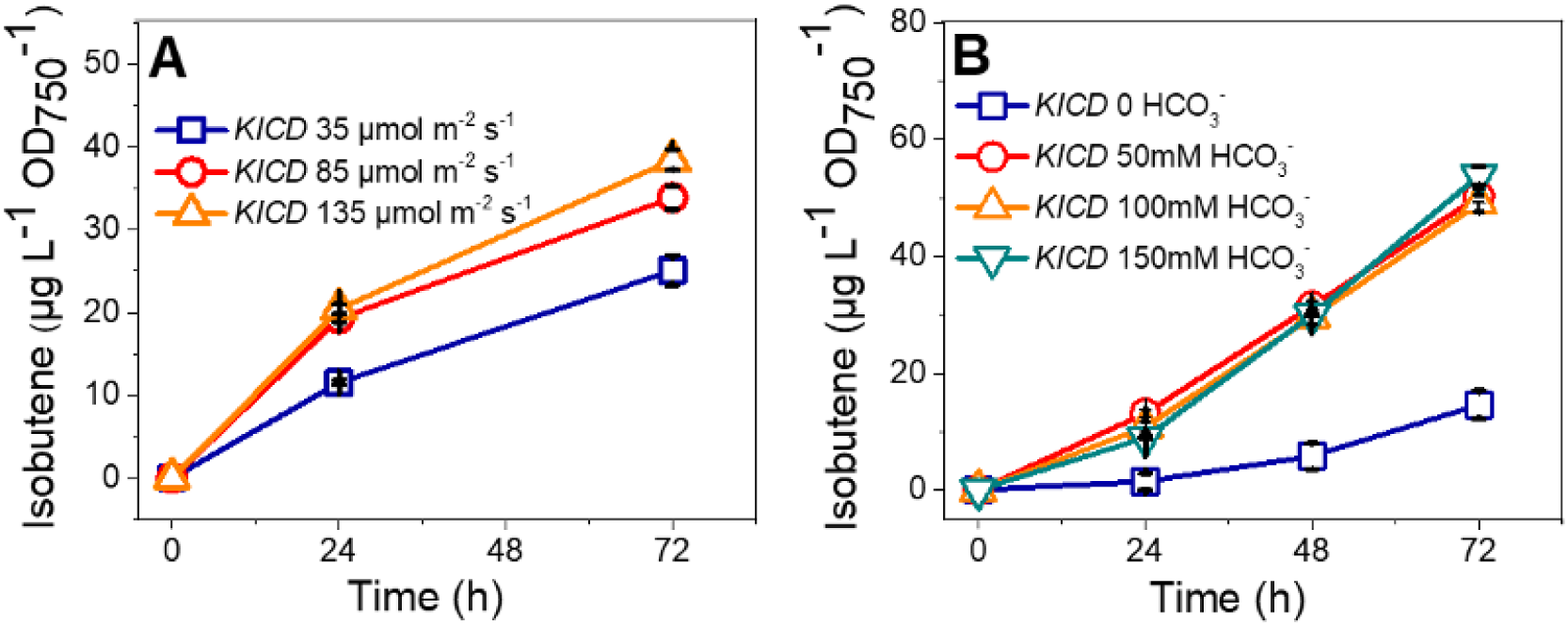
Influence of light intensity and inorganic carbon source on isobutene production in Syn-*Rn*KICD cells. A) Isobutene production was measured in the headspace of cultures cultivated under three light intensities (35, 85, and 135 μmol photons m^-2^ s^-1^). All cultures were uniformly supplemented with 50 mM HCO_3_. B) Isobutene productivity was assessed across a gradient of HCO_3_ concentrations (0 - 150 mM) under 135 μmol photons m^-2^ s^-1^. The cumulative isobutene production was calculated by summing the isobutene produced at 24-h intervals and normalized to each cell culture’s initial OD_750_ values. The initial OD for all cultures was set similarly across all experiments in this study. KICD refers to Syn-*Rn*KICD. Data shown are the average of 3 independent measurements from 3 replicates (±SD).

After 24 h under production conditions, isobutene productivity in Syn-*Rn*KICD cells was observed to be ∼11 μg L⁻¹ OD₇₅₀⁻¹ under LL intensity (Fig. 1A). Higher productivity levels of ∼19 and 20 μg L⁻¹ OD₇₅₀⁻¹ were observed under ML and HL, respectively. These results indicate that, while ML and HL light intensities result in higher isobutene production compared to LL, the difference between ML and HL intensities at 24 h was negligible, and their productivity was proportional to the increased cell density as observed in their OD₇₅₀ (Fig. S1A).

At 72 h, isobutene production reached ∼25 μg L⁻¹ OD₇₅₀⁻¹ under LL, ∼33 μg L⁻¹ OD₇₅₀⁻¹ under ML, and ∼38 μg L⁻¹ OD₇₅₀⁻¹ under HL intensities (Fig. 1A). While HL intensity resulted in the highest isobutene production, it also caused cell bleaching and a decrease in cell density by 72 h (Fig. S1A). In contrast, cells exposed to LL and ML showed no bleaching during this period. This suggests that cell bleaching might result from an imbalance between light energy input and Ci assimilation. Prior studies indicate that cell bleaching can impair the growth, photosynthetic efficiency and productivity of *Synechocystis* (Van Alphen et al., 2018; Vajravel et al., 2020). Adequate nutrient supply, including Ci, can mitigate cell bleaching under such light conditions in cyanobacteria (Markou et al., 2014).

Thus, we hypothesized that excessive light energy relative to available Ci could induce oxidative stress, leading to cell bleaching in *Rn*KICD cells. To test this, cells were exposed to constant HL (135 μmol photons m⁻² s⁻¹) with varying HCO₃⁻ concentrations (0 to 150 mM). The results showed that cells supplemented with higher concentrations of HCO₃⁻ (100 and 150 mM) did not exhibit cell bleaching at 72 h, while those with lower HCO₃⁻ (50 mM) experienced bleaching and reduced growth (Fig. S1B). This suggests that insufficient Ci levels under HL intensity disrupt energy homeostasis, potentially causing oxidative stress and bleaching.

Although isobutene production was observed under all conditions, cells supplemented with HCO₃⁻ exhibited significantly higher growth and isobutene productivity compared to the non-supplemented control cells. The maximum isobutene productivity (∼53 μg L⁻¹ OD₇₅₀⁻¹) was achieved with 150 mM HCO₃⁻. That was nearly 4 times higher than in the control cells (Fig. 1B). Although no significant difference in isobutene productivity was observed between 50 to 150 mM HCO₃⁻ supplementation (49–53 μg L⁻¹ OD₇₅₀⁻¹ at 72 h, Fig. 1B), bleaching occurred earlier in the cultures with low HCO₃⁻ level (50 mM) potentially impacting long-term isobutene production. Notably, control cells (0 mM HCO₃⁻), which showed lower isobutene production (14 μg L⁻¹ OD₇₅₀⁻¹ at 72 h; Fig. 1B), did not exhibit the bleaching phenotype observed in cells with 50 mM HCO₃⁻, nor did they show substantial growth (Fig. S1B). Such production likely originated from stored carbon in the control cells. These results highlight the critical balance of HCO₃⁻ in preventing cell bleaching under HL conditions in Syn-*Rn*KICD cells.

HCO₃⁻ serves as a key substrate for the Calvin cycle, the primary pathway of CO_2_ fixation in cyanobacteria (Bennet, 1991; Miller et al., 1990; Badger and Price 2003). To balance the supply and demand side of photosynthetic metabolism, HL intensities increase the demand for CO_2_ fixation. When HCO₃⁻ levels are limiting, the cell growth is competing with isobutene production pathways. Prior studies also corroborate that excess light can trigger oxidative stress through reactive oxygen species (ROS) overproduction, decreasing photosynthetic efficiency and causing bleaching (Miller and Colman, 1980; Miller et al., 1990; Gordillo et al., 1998; Rodrigues et al., 2023). Our findings emphasize the need to balance light intensity and Ci source availability to maintain *Synechocystis* cell viability and improve isobutene production.

### 3.2. Impact of Nitrogen starvation on the isobutene production in *Synechocystis* cells

#### 3.2.1. Impact of short-term nitrogen starvation on Syn-*Rn*KICD experimental cultures

Previous research by our group implies that KIC serves as substrates for both isobutene and BCAA biosynthesis pathways in *Synechocystis* (Mustila et al., 2021; Schumann et al., 2024). We reasoned that (i) limiting N availability would decrease the metabolic flux toward BCAA synthesis, thus redirecting more carbon toward isobutene production, (ii) the induction of N starvation in *Synechocystis* can provide valuable insights into how N manipulation impacts the partitioning of carbon fluxes between competing biosynthetic pathways in Syn-*RnKICD*. To test these hypotheses, we subjected Syn-*Rn*KICD cells, incubated with 50 mM HCO_3_ under 50 μmol photons m⁻² s⁻¹ light intensity to prevent HL-induced damage, to N starvation and evaluated its effect on isobutene productivity and on major photosynthetic pigments (Fig. 2).

**Figure 2:**
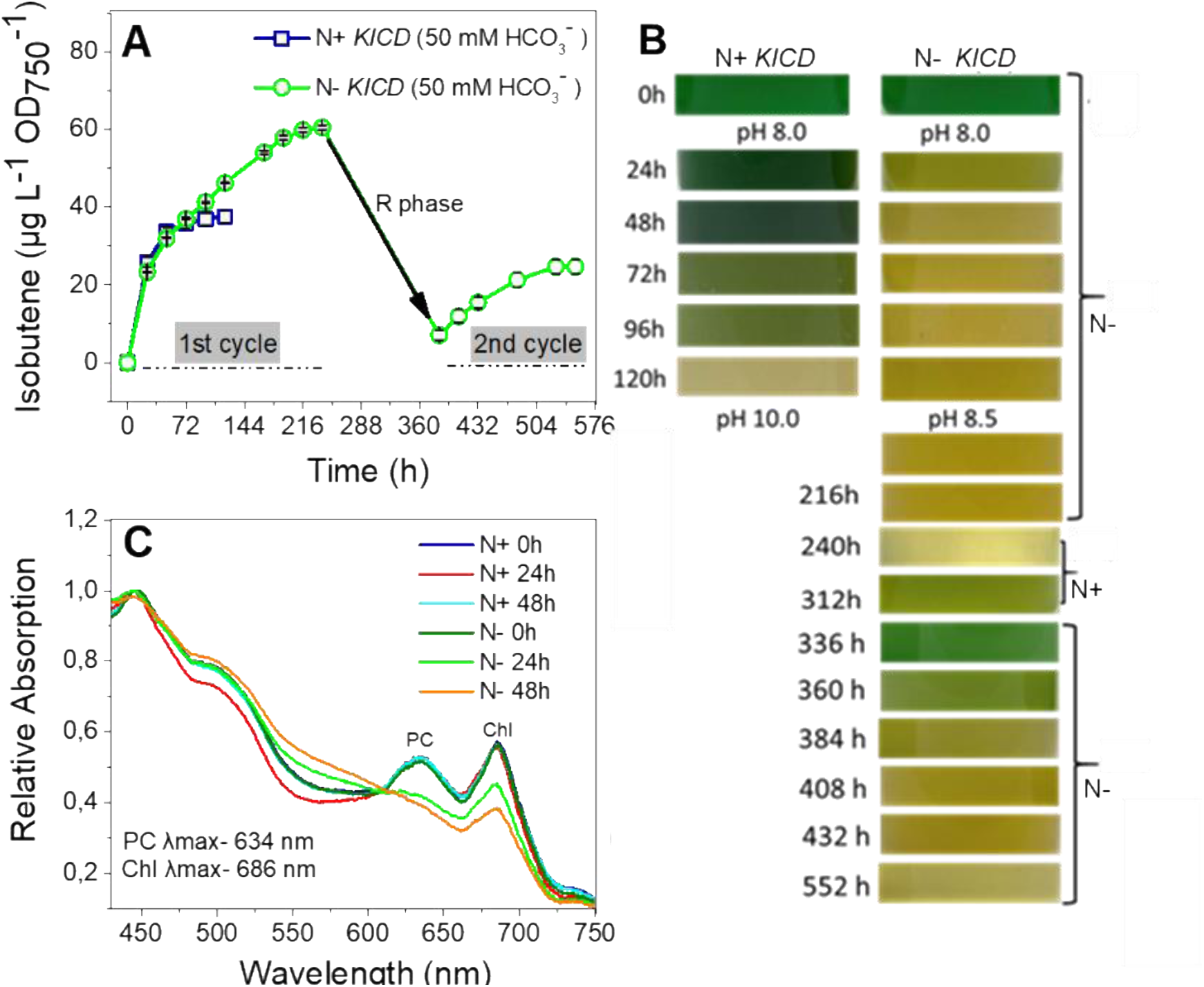
Impact of nitrogen starvation on isobutene production in Syn-*Rn*KICD cells. A) Isobutene productivity was periodically measured in cell cultures supplemented with 18 mM NaNO3 (N+), and without NaNO3 (N-) in BG11 medium. The experimental cultures were cultivated in an appropriate medium for 24 h before being subjected to isobutene production conditions in tightly sealed vials, which were continuously supplied with an appropriate nitrogen-containing medium. The produced isobutene was measured from the headspace of the culture vials every 24 h. B) Phenotypic changes of the cultures were monitored throughout the experiment. C) Comparative whole-cell ABS spectra were obtained to a similar OD750 of the cells which showed phycobilin (λmax = 634 nm) and Chl a (λmax = 686 nm) peaks. The black arrow in Figure A indicates the resuscitation phase, where the reintroduction of NaNO3 into the N- cultures facilitated a recovery process, and a subsequent cycle of N starvation was instituted. All the cultures were maintained under 50 μmol m^-2^ s^-1^ with 50 mM HCO3. Isobutene productivity was normalized to the initial OD750 values of cultures, which was set to similar in all the experimental setups. These results are the average of 3 independent measurements from 3 replicates (±SD). KICD refers to Syn-*Rn*KICD.

We observed that isobutene production occurred regardless of N supply (Fig. 2A). However, N starved cells exhibited chlorosis after 24 h (Fig. 2B), indicating reduced level of major photosynthetic pigments, particularly phycobilins (PB) and Chl (Fig. 2C). The absorption (ABS) spectra results showed that a sharp decrease in PB-to-Chl ratio, a known marker of N deprivation in *Synechocystis,* which was also observed in previous studies in cyanobacteria (Allen and Smith, 1969; Allen et al.,1990). Up to 72 h, there was no significant difference in isobutene production between N supplied (N+) and N starved cells (N-) (Fig. 2A). However, after 96 h, N- cells outperformed N+ cells producing ∼46 μg L⁻¹ OD₇₅₀⁻¹ of isobutene at 120 h, which was ∼1.2 times the amount produced by N+ cells (37 μg L⁻¹ OD₇₅₀⁻¹). At this point, N+ cells showed stagnation in isobutene production (Fig. 2A), along with severe cell bleaching and significant pigments loss (Fig. 2B). On the other hand, N- cells continued to produce isobutene up to 216 h, reaching ∼60 μg L⁻¹ OD₇₅₀⁻¹ (Fig. 2A).

Although both N+ and N- cells had similar isobutene production rates up to 48 h, N- cells outperformed N+ cells from 72 h onward (Fig. S2A). Under the tested conditions, the rate of production was enhanced 10 times as compared to the levels previously reported in *Synechocystis* (Mustila et al., 2021). Initial cell growth, as measured by OD_750_, and Chl content, was 2 to 4 times higher in N+ cells than in N- cells up to 72 h (Fig. S2B and C). At 96 h, N+ cells showed a rapid decrease in their photosynthetic pigments (Fig. S2D) and no pigment peaks were detected by 120 h, likely due to severe cell bleaching which led to no further isobutene production. On the other hand, although N- cells showed reduced pigments levels initially (Fig. 2B), only a gradual decline in the ABS spectra was observed between 48 and 120 h (Fig. S2E) and these N- cells continued to produce isobutene until 216 h (Fig. 2A).

To explore recovery in N- cells, N was reintroduced in the cultures at 252 h. During resuscitation phase (R-phase), these cells regenerated their growth and photosynthetic pigments (Fig. 2B, S2B, C, and F), similar to findings in a previous study on cyanobacteria (Forchhammer & Schwarz, 2019). After initiating N starvation for the second time at 120 h, isobutene production resumed, albeit at lower levels than during the first cycle, producing ∼84 μg L⁻¹ OD₇₅₀⁻¹ of isobutene at 552 h (combining the first and second cycles) (Fig. 2A). This overall productivity is 2.3-fold higher than that of the N+ cells (37 μg L⁻¹ OD₇₅₀⁻¹). These results suggest that N limitation could prolong isobutene production by preventing cell growth and major pigment loss. Furthermore, the recovery after N reintroduction highlights the potential for cyclic N manipulation to optimize long-term isobutene productivity.

A critical question is why N+ cells underwent bleaching and ceased production of isobutene despite being maintained under optimal light conditions (50 μmol photons m⁻² s⁻¹) and with the supply of 50 mM HCO₃. Considering these factors, it is also important to note that cell bleaching occurred later under these conditions (120 h, Fig. 2B) compared to cells exposed to HL (135 μmol photons m⁻² s⁻¹), where cell bleaching was observed as early as 72 h (Fig. S1B). The bleaching in N+ cells may be attributed to Ci limitation probably caused by the competition between overgrowth, and isobutene production in air-tight vials with 50 mM HCO₃⁻ supplementation. In contrast, N+ cells grown in E-flasks did not bleach even after two weeks (data not shown), likely due to the sufficient CO_2_ gas exchange through the cotton- sealed flasks.

Surprisingly, however, N- cells, maintained isobutene production over 552 h even with a reduced amount of photosynthetic pigments (Fig. 2A). In correlation with that several previous studies showed that in cyanobacteria N starvation is initiated with the degradation of nitrogen-rich phycobiliproteins, to supply essential nutrients for cell survival (Allen & Smith, 1969; Spat et al., 2018). Further, N starvation induced chlorosis leads to a dormant state in which cell growth is arrested, and low-level photosynthesis is maintained as a long-term survival mechanism (Forchhammer & Schwarz, 2019; Forchhammer & Selim, 2020). Despite this, CO_2_ fixation can still occur in N starved cells, albeit at lower levels (Allen et al., 1990). In line with these reports, our results showed a significant growth reduction in N- cells compared to N+ cells (Fig. S2B and C). N+ cells exhibited 4.7-fold increase in OD, while N- cells showed a 2-fold increase after 48 h compared to their initial levels (0 h), despite all cultures having similar initial OD values (Fig. S2B and C). Based on this, we suggest that N- cells could still fix CO₂ and are directing carbon toward isobutene production rather than cell growth. These results reveal that the limited growth under N deprived conditions might prolong cell survival and enable isobutene production for a longer period than in N supplied cells.

#### 3.2.2. Impact of prolonged nitrogen starvation on Syn-*Rn*KICD experimental cultures

To determine if further growth restriction in N starved cells could enhance isobutene productivity within a shorter production period, we extended the N starvation period in Syn- *Rn*KICD experimental cultures. Unlike previous experiments where experimental cultures were subjected to 24 h of N starvation (Fig. S2B), the cells in this experiment underwent 72 h of N starvation. In all N starvation experiments, isobutene production in the tightly sealed vials was consistently supplemented with an appropriate nitrogen-containing medium.

As expected, the cells subjected to prolonged N starvation exhibited higher isobutene production at 48 h (29 μg L⁻¹ OD_750_⁻¹), compared to N+ cells (24 μg L⁻¹ OD_750_⁻¹) (Fig. 3A).

**Figure 3:**
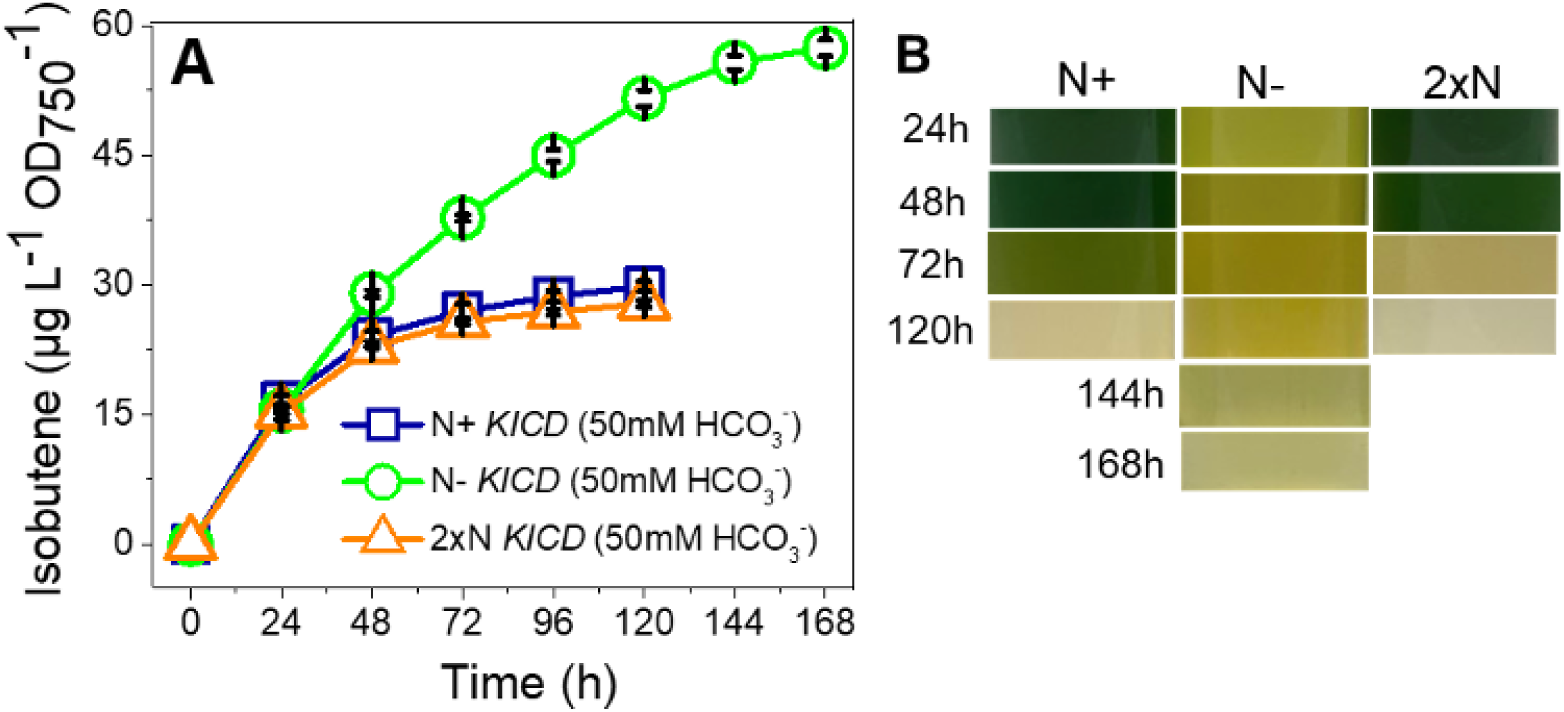
Impact of prolonged nitrogen starvation on isobutene productivity in Syn- *Rn*KICD cells. A) The cumulative isobutene productivity was evaluated in cultures under three distinct conditions: N+, N- and 2xN. All experimental cultures were cultivated in 50 mM HCO_3_ supplemented BG11 medium in the presence (N+ and 2xN) and absence (N-) of NaNO_3_ for 72 h before transferring to tightly sealed vials for determination of isobutene productivity. The cultures for production conditions were continuously supplied with an appropriate nitrogen-containing medium. B) Phenotypic changes of the cultures were monitored throughout the experiment. The cultures were incubated with 50 mM HCO_3_ under 50 μmol photons m^-2^ s^-1^ light intensity. The isobutene productivity was normalized on the initial OD_750_ values of all the cultures. These results are the average of 3 independent measurements from 3 replicates (±SD). KICD refers to Syn-*Rn*KICD.

Furthermore, N- cells showed consistency in their rate of isobutene production over 120 h (Fig. S3A) and OD of these cells slightly increased to1.5 times as compared the initial OD (Fig. S3B and C). In contrast, cells with only 24 h of adaption to N starvation showed a 2-fold increase in OD (Fig. S2B). At 120 h, no pigments were detected in N+ cells, while N- cells still displayed a Chl peak at 686 nm (Fig. S3D and F). Correspondingly, N+ cells showed a decline in isobutene production by 120 h, whereas N- cells continued producing isobutene until 168 h, reaching a productivity of 57 μg L⁻¹ OD₇₅₀⁻¹, nearly twice that of N+ cells (30 μg L⁻¹ OD₇₅₀⁻¹) (Fig. 3A). In the cells adapted to nitrogen starvation for 72 h, maximum isobutene productivity was observed at 168 h (Fig. 3A), while cells with 24 h of N starvation adaptation required 216 h to reach the same level of productivity (Fig. 2A). These results suggest that substantial growth limitation due to the prolonged N starvation can increase isobutene production within a shorter time frame, possibly by redirecting carbon flux toward isobutene production, rather than cell growth.

To assess if severe bleaching in N+ cells was caused by competition for Ci sources between growth and isobutene production, we tested cells with doubled nitrogen content (2xN), assuming that this would enhance initial growth on the expense of isobutene production. We supplemented cultures with 2xN (36 mM NaNO₃) and 50 mM HCO₃. The results showed increased cell growth in cultures supplemented with 2xN by 48 h (Fig. 3B), accompanied by higher OD and Chl levels at 24 and 48 h, respectively, compared to N+ cells (Fig. S3B and C). This increased growth potentially contributed to the marginally lower isobutene production as expected in 2xN cells (28 μg L⁻¹ OD₇₅₀⁻¹) compared to N+ cells (30 μg L⁻¹ OD₇₅₀⁻¹) (Fig. 3A). Further accelerated growth in these 2xN cells led to faster bleaching, which was observed as early as 72 h, compared to N+ cells, where bleaching was observed only at 120 h (Fig. 3B). The ABS spectra also showed that the decrease in photosynthetic pigments was faster in 2xN cells up to 72 h, with no detectable pigments at 96 h (Fig. S3E), whereas N+ cells still showed detectable Chl peaks at that time (Fig. S3D). These findings confirm that the relatively increased growth in 2xN cells resulted in faster pigment degradation, bleaching and relatively lower isobutene production compared to N+ cells, suggesting that competition between cell growth and isobutene production can lead to carbon limitation in the tightly sealed vials over time.

#### 3.2.3. Influence of both bicarbonate and nitrate limitation on isobutene production in Syn-*Rn*KICD cells

To explore whether limiting the cell growth, by reducing Ci availability, would affects bleaching and isobutene production, Syn-*Rn*KIC were cultivated without HCO₃⁻ supplementation. The three culture conditions employed in this experiment were as follows: N+, 2xN, and N-, all without HCO₃⁻. Interestingly, up to 144 h, no noticeable difference in isobutene productivity was observed across all culture conditions in the Ci-starved state (Fig. 4A). N- cells still exhibited chlorosis-induced pigment loss in the absence of HCO₃⁻ (Fig. 4B). Up to 48 h, no growth was observed in the cells under the 2xN condition (Fig. S4A).

**Figure 4:**
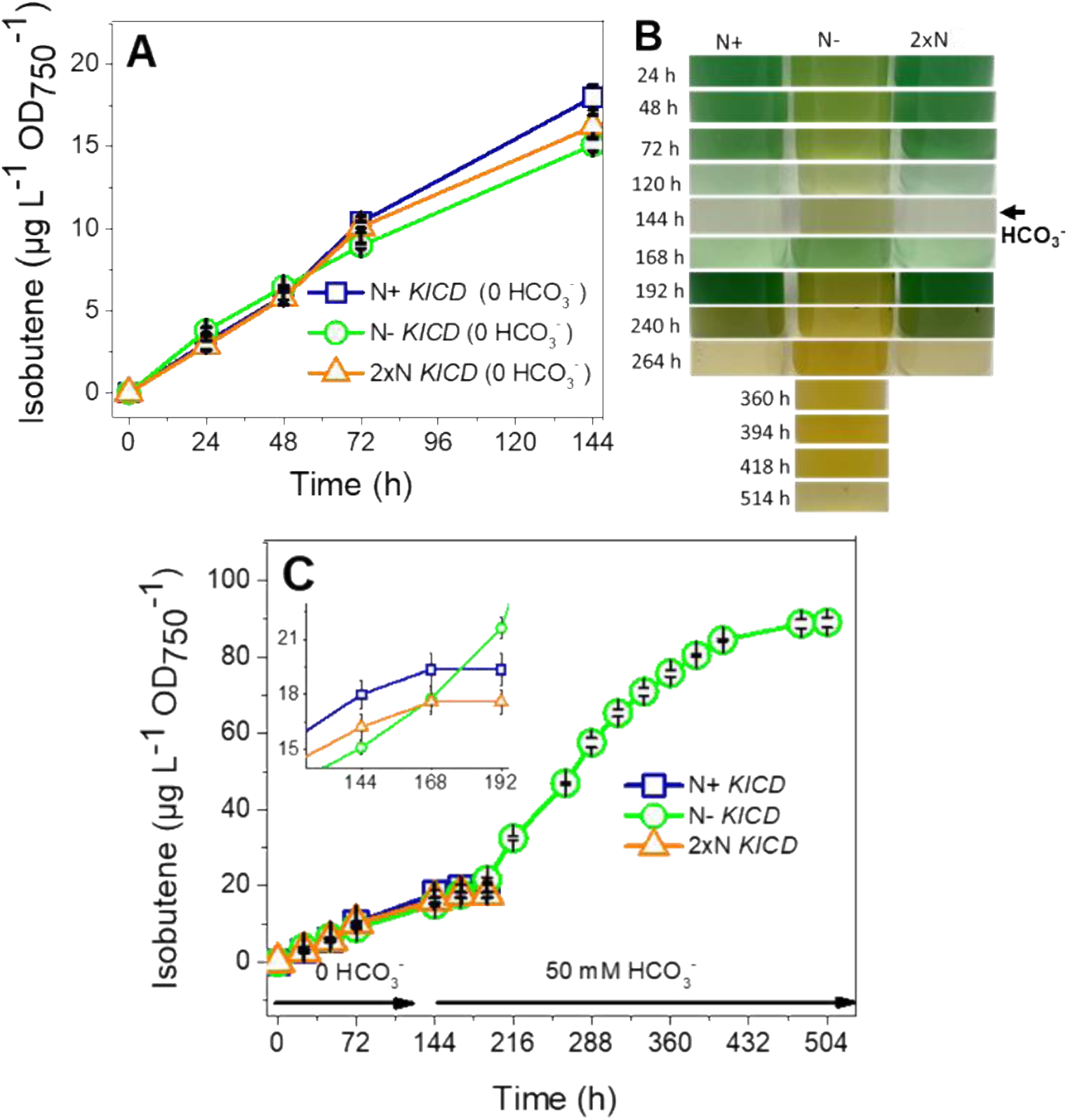
Impact of nitrogen and carbon limitation on isobutene production in Syn- *Rn*KICD cells. A) The isobutene productivity was monitored in Syn-*Rn*KICD cells under three distinct conditions: N+, N-, and 2xN without addition of HCO_3_. The cultures were adapted to the different nitrogen concentrations for 24 h before transferring to tightly sealed vials for determination of isobutene productivity. The cultures for production conditions were continuously supplied with an appropriate nitrogen-containing medium. B) Phenotypic changes of the cultures were monitored throughout the experiment. C) Cumulative isobutene production was examined under: N+, N-, and 2xN and 50 mM HCO_3_ was added after 144 h of production period.The cultures were illuminated with 50 μmol photons m⁻² s⁻¹. The isobutene productivity was normalized based on the initial OD_750_ values of all the cultures. These results are the average of 3 independent measurements from 3 replicates (±SD). KICD refers to Syn- *Rn*KICD.

In corresponding with that, a significant decrease of photosynthetic pigments was observed in N+ and 2xN cells after 96 h (Fig. S4B and C). In contrast, N- cells did not show a decrease in pigment content (Fig. S4D). Interestingly, for an extended period (up to 144 h) all cells continued to produce almost similar amount of isobutene but at a low level. This suggest that that in the absence of HCO₃⁻ no significant difference in isobutene productivity was observed between N supplied (N+ and 2xN) and N starved (N-) cells (Fig. 4A). However, it is important to note that without HCO₃⁻, overall isobutene productivity dropped significantly to 10 and 9 μg L⁻¹ OD_750_⁻¹ at 72 h (Fig. 4A), as compared to 35 and 38 μg L⁻¹ OD_750_⁻¹ with HCO₃⁻ supplementation in N+ and N- cells, respectively (Fig. 2A). These results suggest that the differences in isobutene production between N+ and N- cells were only observed in the presence of HCO₃⁻.

To investigate whether the re-addition of HCO₃⁻ could result in any differences in the regeneration of cell growth and isobutene production among these Ci-deprived cells, HCO₃⁻ was added to all cultures (N+, 2xN, and N-) at 144 h. Surprisingly, all cultures exhibited growth, as indicated by phenotypic changes observed at 192 h (Fig. 4B). However, increased isobutene production was only observed in N- cells. In contrast, no significant increase in the isobutene production was noticed in N supplied cells (N+ and 2xN), which, on the other hand, show a substantial increase in their cell growth (Fig. 4B), probably leading to Ci depletion, thus, the ability to produce isobutene in these N supplied cells might be diminished. Surprisingly, N deprived cells continued producing isobutene for 480 h, reaching ∼89 μg L⁻¹ OD_750_⁻¹. These results indicate that CO₂ fixation using HCO_3_ as Ci source is most likely more directed toward isobutene production since cell growth is restricted in N deprived cells. Furthermore, the observed difference in isobutene production between N supplied and N deprived cells clearly to be primarily due to differences in their cell growth, which were highly influenced by the addition of HCO₃.

Under N limitation, growth is reduced, likely at least partly due to decreased BCAA biosynthesis. Such limited growth might lead to metabolic redirection of carbon toward isobutene production. As a result, isobutene formation becomes less dependent on growth, with carbon flux shifting away from BCAA biosynthesis toward isobutene production.

#### 3.2.4. Assessment of isobutene production in the Syn-F336V strain under nitrogen starvation

In a previous report, we showed that by introducing an engineered *Rn*KICD, with enhanced substrate specificity, to *Synechocystis* (strain Syn-F336V) an increased *in vivo* isobutene production was achieved (Schumann et al., 2024). Here, we examine the effects of N starvation on isobutene production in the Syn-F336V cells to determine if the earlier observed results were specific to the unmodified *Rn*KICD enzyme or represent a broader metabolic shift. Further, we expect Syn-F336V cells to reach higher production level of isobutene than Syn- *Rn*KICD under N limitation. This expectation stems from two key factors: the reduced BCAA synthesis under N limitation, which redirects more carbon precursors toward isobutene production, and the F336V’s improved ability use KIC over HPP as substrate. This approach can help to clarify how both enzyme and cultivation optimization can synergistically increase isobutene yields.

Our results showed that the N supplied Syn-F336V cells exhibited a 1.6-fold increase in isobutene production, reaching 60 µg L⁻¹ OD_750_⁻¹ at 96 h (Fig. 5A), as compared to the 37 µg L⁻¹ OD_750_⁻¹ by Syn-*Rn*KICD cells (Fig. 2A) under identical conditions. Thus, confirming that Syn-F336V cells are significantly more efficient at producing isobutene than the Syn- *Rn*KICD cells. When subjected to N starvation, the Syn-F336V cells showed even higher isobutene production, reaching ∼80 µg L⁻¹ OD_750_⁻¹ at 96 h, surpassing the productivity of their N supplied cells (Fig. 5A). N+ Syn-F336V cells, like N+ Syn-*Rn*KICD cells, displayed increased cell growth up to 72 h, followed by cell bleaching at 96 h (Fig. 2B and 5B), with a decline in their isobutene production. Further, under N- conditions, Syn-F336V cells exhibited chlorosis, a typical symptom of N starved cyanobacterial cells. Despite this, the isobutene production continued to increase for 72 h in Syn- F336V N- cells, reaching a maximum yield of ∼120 µg L⁻¹ OD_750_⁻¹ at 216 h (Fig. 5A). The peak of isobutene productivity in the Syn- F336V N- cells was 2-fold higher than in the Syn-*Rn*KICD N- cells over the same period (Fig. 2A and 5A). These results suggest that the Syn-F336V cells benefit from reduced BCAA biosynthesis under N limitation and increased substrate specificity of the *Rn*KICD enzyme towards KIC, leading to overall enhanced isobutene production than Syn-*Rn*KICD. Both Syn- F336V and Syn-*Rn*KICD strains showed increased isobutene production under N limitation, suggesting that this response is specific to the synthetic isobutene production pathway in *Synechocystis* cells. In this pathway, branched-chain amino acid (BCAA) biosynthesis and isobutene production compete for the same substrate, KIC (Fig.11).

**Figure 5:**
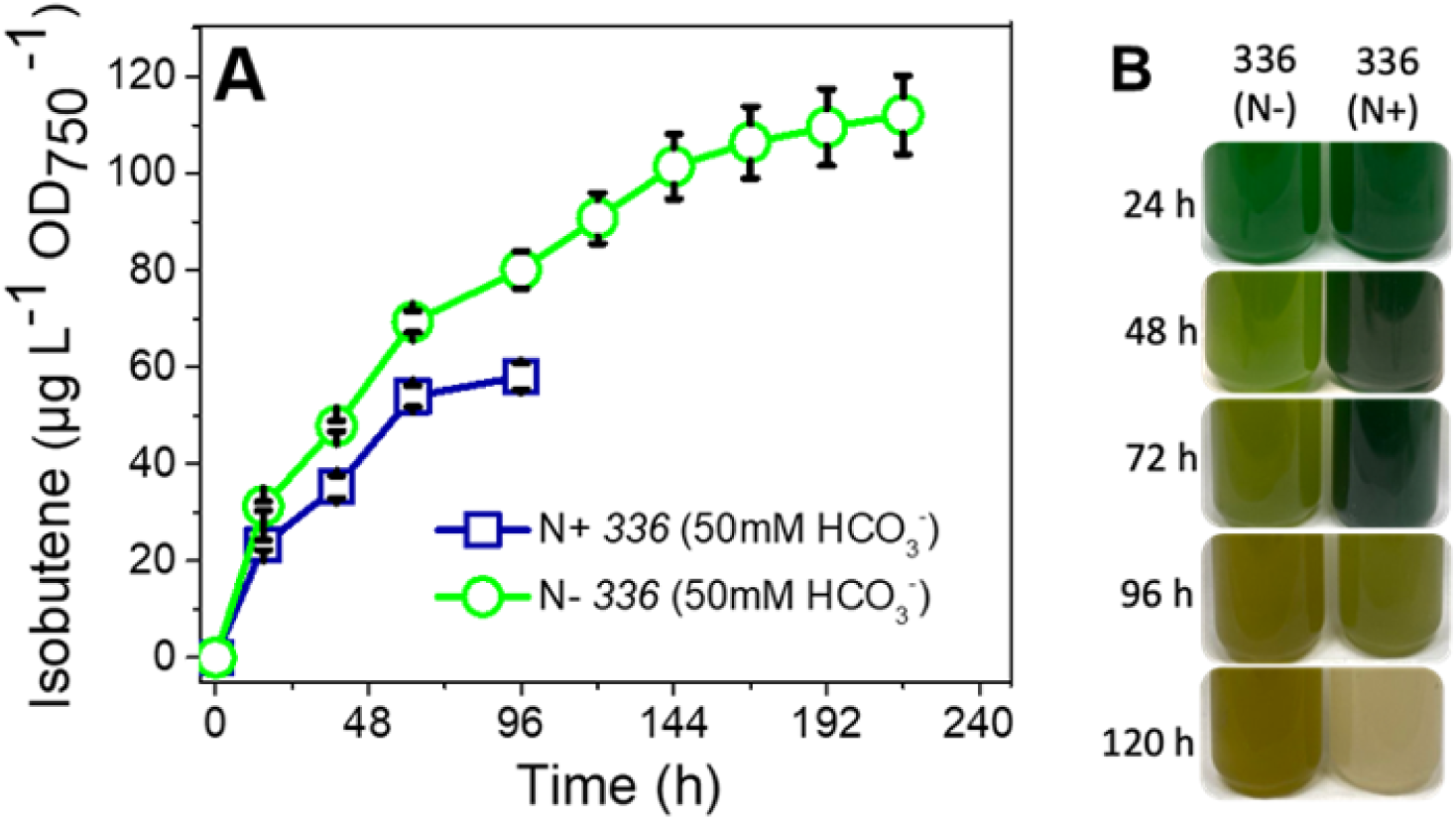
Impact of nitrogen starvation on isobutene production in Syn-F336V cells. A) The cumulative productivity of isobutene was determined in the cells incubated in BG11 medium with and without supplemented NaNO₃. B) Phenotypic changes of the cultures were monitored every 24 h throughout the experiment. The cultures were illuminated with 50 μmol photons m^-2^ s^-1^ and supplemented with 50 mM HCO₃. The isobutene productivity was normalized to the initial OD_750_ values of all the cultures. These results are the average of 3 independent measurements from 3 replicates (±SD). 336 refers to Syn-F336V.

Taken together the results suggest that when N+ cells are supplied with HCO₃⁻ a competition for carbon may occur between their cell growth and isobutene production. This competition will eventually lead to carbon limitation, inducing metabolic stress and cell damage in closed culture vials, as shown in Figures 2 to 5. Conversely, N starvation-induced growth arrest in N- cells could direct the supplied Ci source primarily towards isobutene production for an extended period of time, as demonstrated in Figures 2 to 5. Although N starved cells exhibited prolonged isobutene production as compared to N+ cells under HCO₃⁻- supplementation (Figs. 2-5), it is essential to note that by time these N deprived cells experienced significant nutrient stress, leading to substantial loss of major photosynthetic pigments (Figs. S2 and S3) and led to a halted isobutene production. Previous studies have shown that while N starved cells can sustain low levels of photosynthesis (Sauer et al., 2001) and CO₂ fixation (Allen et al.,1990), this compromised photosynthetic capacity may ultimately limit long-term cell survival. Thus, utilizing N starved cells may not be ideal for sustainably maximizing isobutene production. Based on these insights, we propose that long-term improvements in isobutene production can be achieved by adopting strategies such as physically restricting cell growth while ensuring the provision of optimal nitrogen and carbon sources to support robust metabolic function.

### 3.3. Entrapment of Syn-*Rn*KICD cells in PVA hydrogel beads

In this section, we employed a cell entrapment strategy for prolonged cell viability and enhanced isobutene production in Syn-*Rn*KICD and Syn-F336V cells by limiting cell growth under the controlled availability of Ci and N sources. For this approach, we used 6% PVA hydrogel crosslinked with 4% H_3_BO_3_ to create a entrapped cell beads. The rationale behind this approach is that physically entrapping cyanobacterial cells can limit their growth, redirecting energy and carbon toward product formation rather than biomass accumulation, as supported by previous studies (Kosourov et al., 2014; Seibert et al., 2018; Vajravel et al., 2020). During the droplet-based bead formation process, using a 5:6 ratio of cell wet mass to PVA hydrogel, we encountered a challenge: the PVA hydrogel tended to adhere, preventing the formation of distinct beads in the presence of the crosslinking agent B(OH)₄⁻. This issue might have resulted from insufficient electrostatic repulsion or steric stabilization among the PVA hydrogel particles, leading to clumping (Wu & Wisecarver et al 1992). To address this, we incorporated a small amount of SA (0.5%) into the PVA hydrogel mix. Because SA can facilitate bead formation due to its ability to form crosslinked structures via ionic interactions, enhancing stabilization. Distinct hydrogel beads were promptly generated following this modification: the cell-PVA-SA mixture, at a 5:6:0.5 ratio, was introduced into a solution of crosslinking agents, H₃BO₃ (4%) and CaCl₂ (2%).

We first evaluated the effects of entrapping Syn-*Rn*KICD cells in PVA hydrogel beads’ stability in the presence of HCO_3._ After crosslinking duration of 5 h at 30 °C, the beads initially appeared stable, but, upon the addition of HCO₃⁻, bead-disintegration was observed at 24 h, resulting in a cell suspension (Fig. 6A). The shorter crosslinking duration time may have contributed to the early disintegration of the beads. Thus, we extended the crosslinking duration from 5 h to 20 h without changing the temperature, aiming to strengthen the interactions between the hydrogel and crosslinking agents, thereby creating more stable hydrogel beads. Although the extended crosslinking time (20 h) showed an increased stability of PVA entrapped cell beads initially in presence of HCO_3_, the beads began to disintegrate after 96 h of HCO₃^-^ supplementation (Fig. 6B). These results suggest that while longer crosslinking period can postpone bead breakdown in the presence of HCO₃, it cannot completely prevent it.

**Figure 6:**
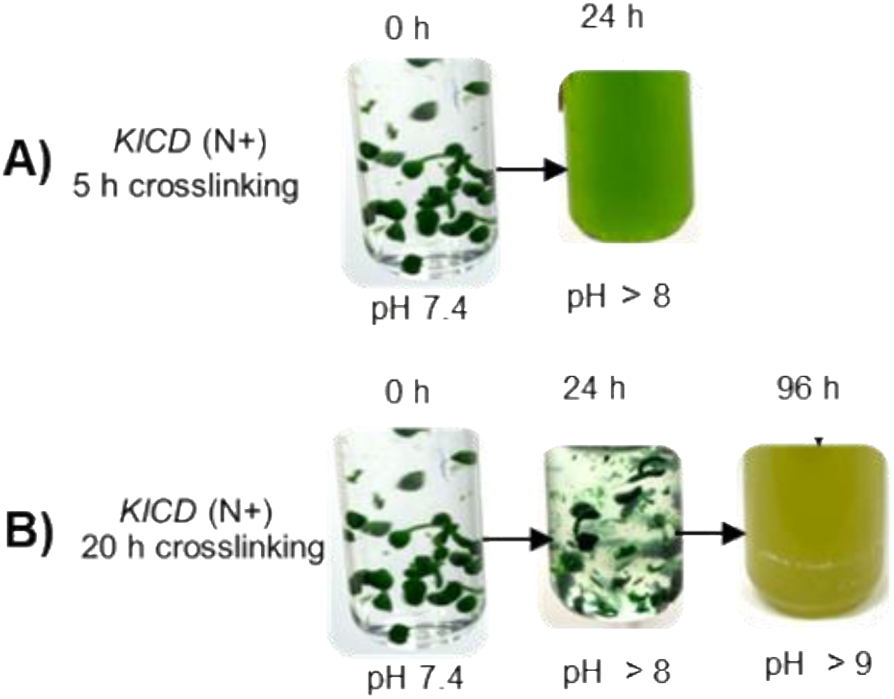
Impact of crosslinking duration on the stability of PVA-SA entrapped Syn- *Rn*KICD cell beads. A) After 5 h of crosslinking, distinct beads were observed, but they disintegrated within 24 h after the addition of 50 mM HCO₃⁻. B) With a crosslinking duration of 20 h, bead stability was maintained even after the addition of HCO₃⁻ for up to 96 h. A 5:6:0.5 ratio of cell wet mass: PVA: SA was employed for droplet-based entrapped cell beads using 4% H_3_BO_3_ and 2% CaCl_2_ crosslinking agents. the entrapped cells were refreshed with BG11 medium, transferred into tightly sealed vials, illuminated with 50 μmol photons m^-2^ s^-1^. KICD refers to Syn-*Rn*KICD.

We also observed a significant increase in pH, from ∼ 7.4 to 9, in the entrapped cell cultures. Despite initially supplementing the entrapped cells with a buffered medium, the addition of 50 mM HCO₃ likely contributed to the pH rise. An increase in alkaline pH can destabilize PVA- SA beads crosslinked by B(OH)₄⁻ and Ca²⁺ ions by altering the ionization states, weakening crosslink density, and potentially causing excessive swelling or hydrolysis. This destabilization may have led to bead disintegration and the release of free cells into the medium.

#### 3.3.1. Effect of pH on the stability of PVA-SA entrapped Syn-*Rn*KICD beads

Our previous results indicated that the stability of PVA-SA hydrogel beads may be compromised under alkaline conditions (Fig. 6). Although *Synechocystis* typically thrives in a pH range of 7.0 to 9.0, maintaining a neutral pH (∼7) could be essential for preventing bead disintegration and ensuring cell viability. By adjusting the medium’s pH to neutral every 24 h, we observed that the hydrogel beads remained stable without disintegration, even after 96 h of incubation with HCO₃⁻ (Fig. 7A). These results underscore the critical importance of maintaining a neutral pH for the stability of entrapped cells in the PVA-SA hydrogel beads. Next, we investigated the isobutene-producing capacity of PVA-SA entrapped Syn-*Rn*KICD cells. 120 h after bead formation, the entrapped Syn-*Rn*KICD cells were transferred into tightly sealed vials and placed under illumination at 50 μmol m⁻² s⁻¹ (post adaptation conditions). We monitored isobutene productivity in the headspace of the vials at 24 h intervals. Remarkably, isobutene production was observed at 144 h after bead formation (24 h of post-adaptation conditions), as shown in Fig. 7B. The isobutene productivity reached ∼40 mg/g DW after 216 h of post-adaptation under continuous illumination (336 h after bead formation).

**Figure 7:**
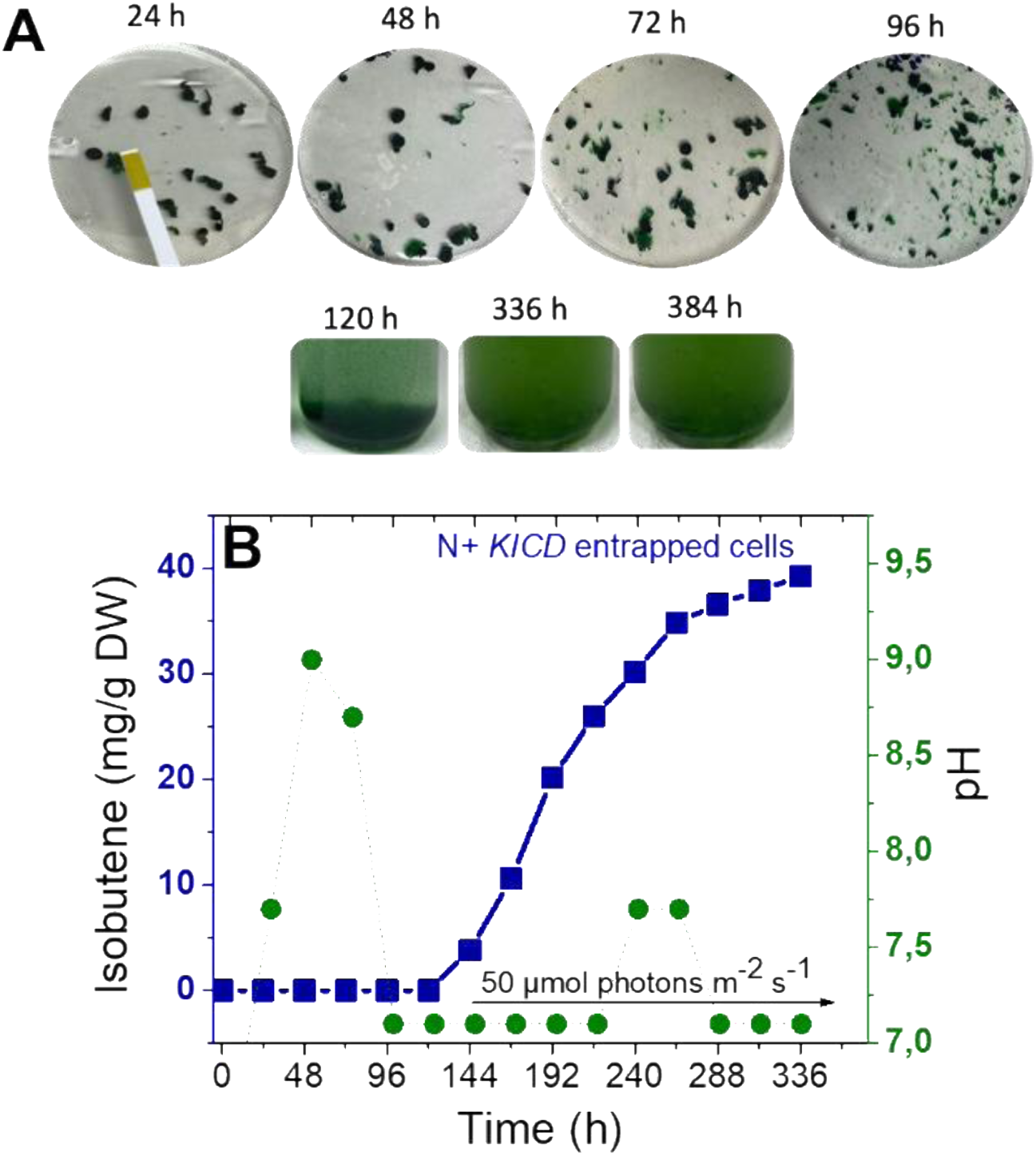
**Impact of neutral pH on the stability of PVA-SA-entrapped Syn-*Rn*KICD cell beads**. A) The phenotypic changes of the entrapped cell beads were observed in the presence of 50 mM HCO₃, with the pH level adjusted to ∼7 – 7.4 with 5N HCl every 24 h. B) Entrapped Syn-*Rn*KICD cells were transferred to sealed vials at 120 h after bead formation and illuminated at 50 μmol photons m⁻² s⁻¹ (post-adaptation conditions). Isobutene production was detected in the vial headspace at 144 h (24 h of post-adaptation). The cumulative isobutene production was monitored for 336 h. The cell entrapment was performed using the following formulation: 5:6:0.5 wet cell mass:PVA:SA and crosslinked with 4% H_3_BO_3_ and 2% CaCl₂ for 20 h. After 20 h of crosslinking duration, the entrapped cells were washed with BG11 medium with NaNO_3_, illuminated with 50 μmol photons m^-2^ s^-1^. The productivity of isobutene was normalized on the initial DW of the cells. The pH level adjusted to ∼7 – 7.4 every 24 h throughout the experiment. KICD refers to Syn-*Rn*KICD.

This observation is particularly notable when compared to the suspension cultures, which exhibited rapid bleaching and cessation of isobutene production within 120 h (Fig. 2). In contrast, entrapped cells within the PVA-SA hydrogel beads did not show any cell bleaching, even after 216 h of incubation in tightly sealed vials, as evidenced by their phenotypic morphology (Fig. 7A). However, after 336 h, partial bead disintegration was observed, but the cells did not exhibit any bleaching phenotype. Such non-bleaching cells likely contributed to the extended period of isobutene production by the entrapped cells (Fig. 7B). These results demonstrate that maintaining a neutral pH in the entrapped cell cultures can enhance the structural integrity of the PVA-SA hydrogel beads for a longer period, which, in turn, could prevent excess growth, cell bleaching and promotes long-term isobutene production. However, despite maintaining a neutral pH over an extended period, the beads showed disintegration and cell suspensions were observed, and distinct beads were no longer visually detectable after 336 h (Fig. 7A). The disintegration of the beads could be attributed to excessive cell growth over time under the supplied nitrogen and HCO₃⁻ condition. This highlights the eventual limitations of hydrogel bead stability and emphasizes the need to optimize the cell-to-hydrogel ratio to strictly limit cell growth, thereby enhancing bead stability.

#### 3.3.2. Impact of doubling hydrogel-to-cell ratio on the stability of PVA-SA entrapped cell beads and isobutene productivity

In our continued efforts to improve the stability of PVA-SA-entrapped cell beads and enhance isobutene productivity, we focused on the impact of increasing the hydrogel-to-biomass ratio. Considering that denser hydrogel may further restrict cell expansion and proliferation, we adjusted the cell wet mass to hydrogel ratio (cell:PVA:SA) from 5:6:0.5 to 5:12:1, while maintaining a neutral pH throughout the experiment. This experiment was designed to investigate how an increase in the hydrogel-to-biomass ratio, combined with neutral pH adjustments, influences PVA-SA-entrapped cell beads’ stability and isobutene productivity in the presence of HCO₃⁻ supplementations. Additionally, we examined the entrapped cells under both N supplied and N deprived conditions. This was motivated by our earlier experiment, which showed that nitrogen-supplied cell suspensions exhibited significant cell growth but decreased isobutene production, whereas nitrogen-deprived cells limited growth and increased isobutene production. Such comparison between N+ and N- conditions can help us better understand the different growth and production dynamics of PVA-SA-entrapped cells. Before bead formation, Syn-*Rn*KICD experimental cells were acclimated to N+ and N- cultivation conditions. The 20-hour crosslinking phase resulted in distinct and stable bead formation, as evidenced by their phenotypic morphology (Fig. S5). Isobutene production in the entrapped cells was then monitored at 24 h intervals in the headspace of tightly sealed vials.

The results showed no significant difference in isobutene productivity between N+ and N- entrapped cells up to 72 h (Fig. 8A). However, at 96 h, N- cells exhibited higher isobutene production, reaching ∼28 mg/g DW, while N+ cells produced ∼19 mg/g DW. Although the beads remained stable up to 144 h under both N+ and N- conditions, at 192 h partial beads’ disintegration was observed under N+ conditions, as shown by their phenotypic morphology (Fig. 8B). However, the rate of isobutene production continued to be monitored under both conditions but at a lower level. At 216 h, 25 mM HCO₃⁻ was added to test if the isobutene production could be further increased. This addition slightly improved isobutene production in both N+ and N- cells from 264 h onwards.

**Figure 8:**
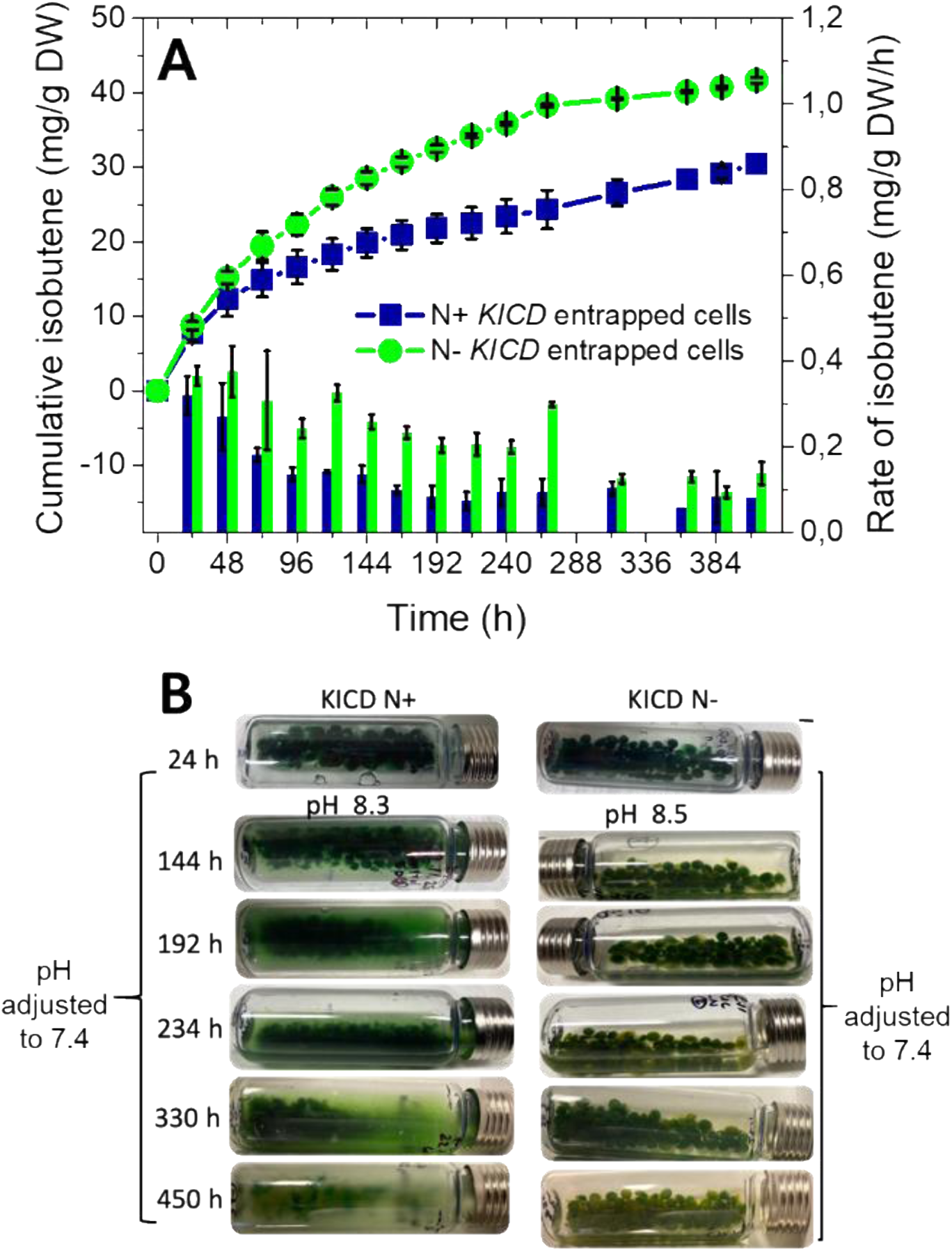
Enhanced bead stability in Syn-*Rn*KICD cells with a 2-fold higher PVA-SA hydrogel-entrapped cell system under N- conditions. A) The cumulative productivity (lines) and rate of isobutene (bars) were monitored in the entrapped cells. B) The physical characteristics of the entrapped cell beads were observed throughout the experiment. The hydrogel beads were prepared with a 5:12:1 ratio of cell wet mass: PVA:SA using 4% H_3_BO_3_ and 2% CaCl₂ crosslinking. The pH of the cultures was monitored every 24 h and adjusted to neutral ∼7.4. Isobutene production was determined in entrapped cells under N+ and N- culture conditions, illuminated with 50 μmol photons m^-2^ s^-1^, and supplemented with 50 mM HCO₃ at day 1, and additional 25 mM HCO₃ was provided at 216 h. The isobutene productivity was normalized on the initial DW of the cells used for entrapment. These results are the average of 3 independent measurements from 3 replicates (±SD).

At 412 h, the N- entrapped cells still exhibited stable beads and higher isobutene productivity, reaching ∼41 mg/g DW, compared to N+ entrapped cells (∼30 mg/g DW) (Fig. 8). No bead disintegration was observed in N- cells at this point (Fig. 8B), suggesting that the combination of N limitation and entrapment-mediated cell growth limitation did not compromise bead stability, cell viability and isobutene production. Further under N- conditions, the entrapped cells not only prolonged their isobutene production but also showed delayed chlorosis response up to 412 h (Fig. 8B), whereas N starved cell suspensions exhibited chlorosis within 24 h (Fig. 2B). These observations indicate that entrapping cells in PVA-SA hydrogel beads could further enhance cell viability and isobutene production for an extended period under N deprived conditions.

On the other hand, bead disintegration was clearly visible under N supplied conditions starting at 192 h, with gradual leakage of cells into the medium, as evidenced in Fig. 8B. Despite this disintegration, isobutene production from the entrapped N+ cells continued for up to 412 h (Fig. 8A), significantly longer than in suspension cells, which only produced isobutene up to 120 h (Fig. 2A). However, the disintegration of beads eventually led to cell suspension and subsequent cell bleaching by 450 h (Fig. 8B). This disintegration of beads could be due to excessive cell growth, as N+ cells tend to exhibit strong growth responses in the presence of HCO₃⁻ supplementation, as seen in the suspension cells (Fig. 2A). These results indicate that while the 2-fold increase in hydrogel-to-cell ratio and maintaining neutral pH prolonged bead stability, the cells under N supplied conditions still experienced delayed bleaching due to delayed beads’ disintegration to cell suspension. Therefore, further optimization of the cell entrapment procedure is needed, particularly for cells under N supplementation, to achieve prolonged isobutene production in this case.

#### 3.3.3. Impact of a sixfold increase in hydrogel-to-cell ratio on the stability of PVA-SA entrapped cell beads and isobutene productivity

Though increasing 2x higher hydrogel to biomass improved beads’ stability for a certain period, our attempts to effectively limit cell growth and prolong isobutene production under N and HCO_3_ conditions were less effective, as seen in bead disintegration, cell leakage, gradual bleaching, and relatively lower isobutene production (Fig. 8). To address this issue, we further experimented with a six-fold increase in the hydrogel-to-cell ratio, changing it from 5:12:1 to 5:36:3 (cell:PVA:SA), aiming to limit cell growth and prevent bead disintegration. This modification in the cell-to-hydrogel ratio led to visually stable and distinct beads for Syn- *Rn*KICD and Syn-F336V cells (Fig. 9).

**Figure 9:**
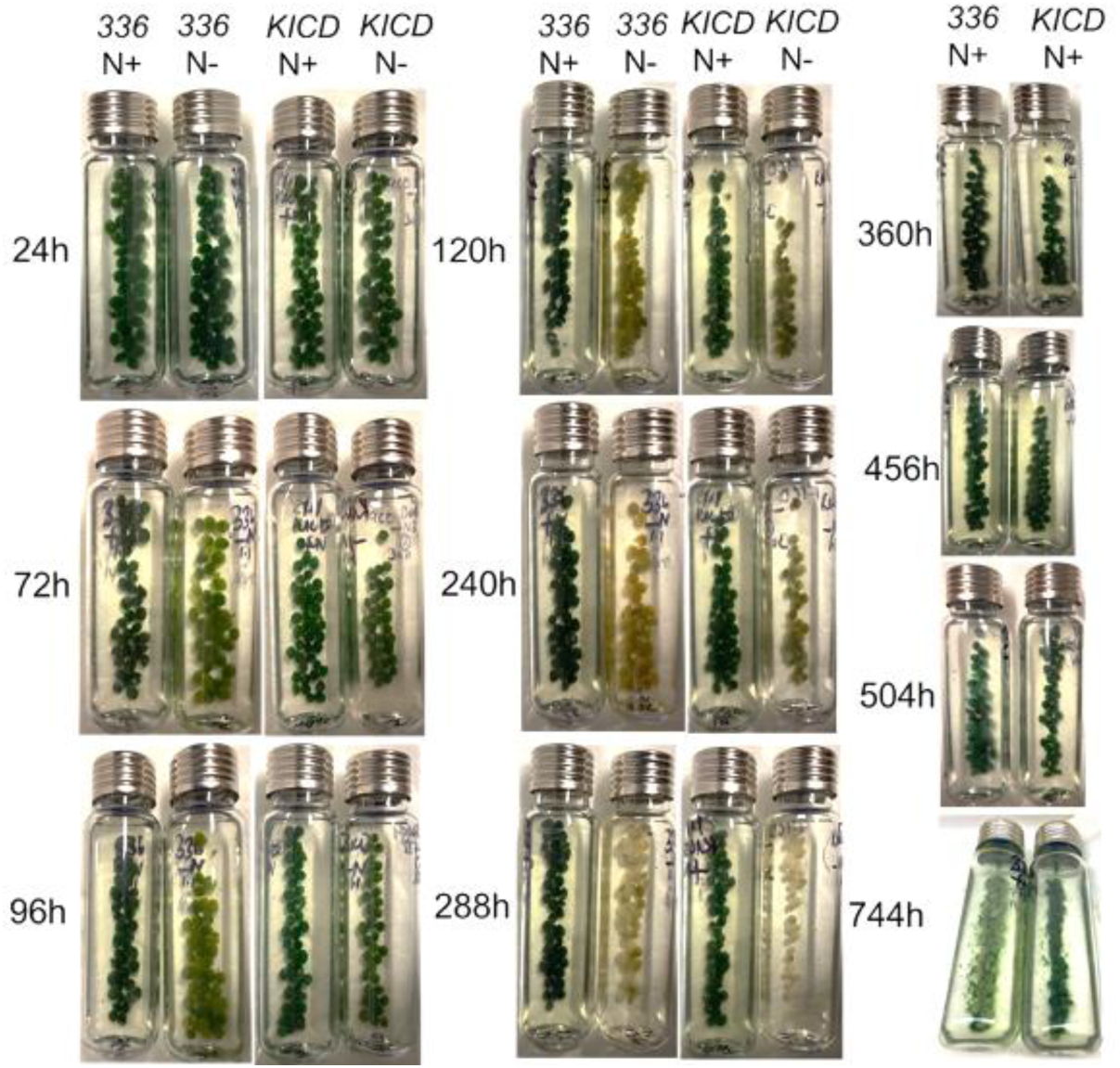
Extended bead stability in Syn-F336V and Syn-*Rn*KICD cells with a 6-fold higher PVA-SA hydrogel-entrapped cell system under N+ and N- conditions. The hydrogel beads were prepared with the following ratio between the cell wet mass: PVA: SA was 5:36:3 using H_3_BO_3_ and CaCl₂ crosslinking. pH was adjusted to neutral throughout the experiments. The entrapped cells were illuminated under 50 μmol photons m^-2^ s^-1^ with addition of 50 mM HCO₃. A similar amount of initial biomass was used across all cultures. 336 refers to Syn- F336V, and KICD refers to Syn-*Rn*KICD.

Up to 196 h, there was no significant difference in isobutene production observed in the entrapped Syn F336V cell beads under N supplied and N starved conditions (Fig. 10A). Interestingly, at 216 h, a notable increase in isobutene production (∼67 mg/g DW) was observed in Syn-F336V N+ cells. Conversely, no further increase was observed in the N- cells, which reached a maximum productivity of ∼57 mg/g DW. On the other hand, Syn-F336V N+ cells continued to increase their isobutene production, achieving ∼91 mg/g DW at 408 h. A similar trend was observed in entrapped Syn-*Rn*KICD cells under the same production conditions, with only slight differences in isobutene productivity between N+ (∼38 mg/g DW) and N- (∼45 mg/g DW) cells up to 216 h (Fig. 10B). After 216 h, no further increase in productivity was observed in N- cells. At this point, a significant increase in isobutene production was observed in N+ Syn-*Rn*KICD cells, which was continuing to rise until 408 h and reached ∼54 mg/g DW. These results showed that both Syn-F336V and Syn-*Rn*KICD strains exhibited higher isobutene production under N supplied conditions than N starved conditions after 216 h (Fig. 10A and B).

**Figure 10:**
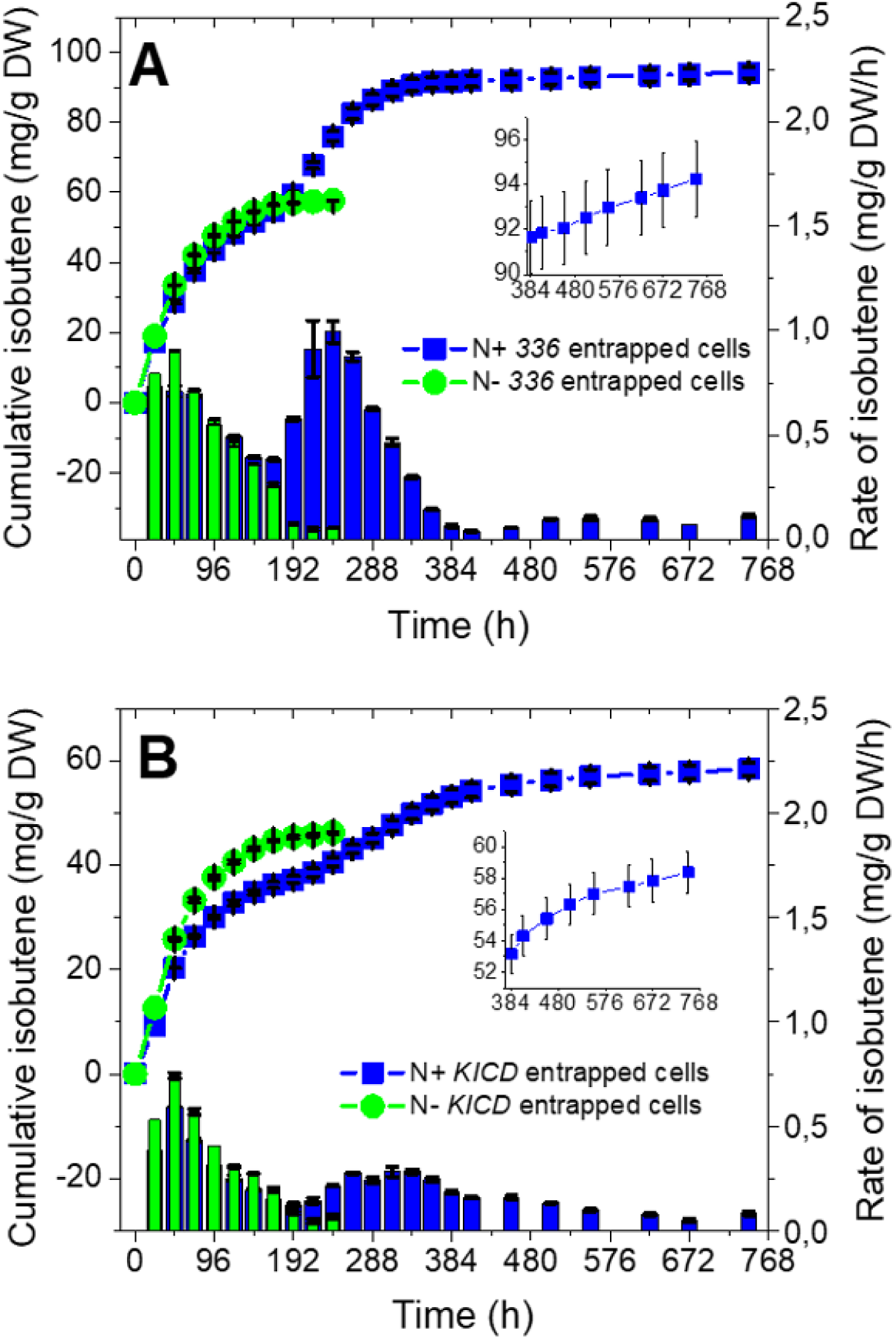
Extended bead stability and isobutene productivity in Syn-F336V and Syn-*Rn*KICD cells with a 6-fold higher PVA-SA hydrogel-entrapped cell system under N+conditions. The cumulative productivity (lines) and rate of isobutene (bars) in the entrapped A) Syn-F336V and B) Syn-*Rn*KICD cells under N+ and N- culture conditions. The beads were prepared with the ratio: 5:36:3 for the wet cell weight: PVA: SA, crosslinked with H3BO3 and CaCl₂. pH was adjusted to neutral throughout the experiments. A similar amount of initial cell mass was used across all cultures. The productivity of isobutene was normalized to the initial DW of the cells. Inlets in Figures A and B showed an enlarged view of the prolonged production of isobutene. The cells were illuminated with 50 μmol photons m⁻² s⁻¹ with addition 50 mM HCO₃ at the start of each experiment, additional 25 mM NaHCO₃ was added at 408 h. These results are the average of 3 independent measurements from 3 replicates (±SD). 336 refers to Syn-F336V, and KICD to Syn-*Rn*KICD.

**Figure 11:**
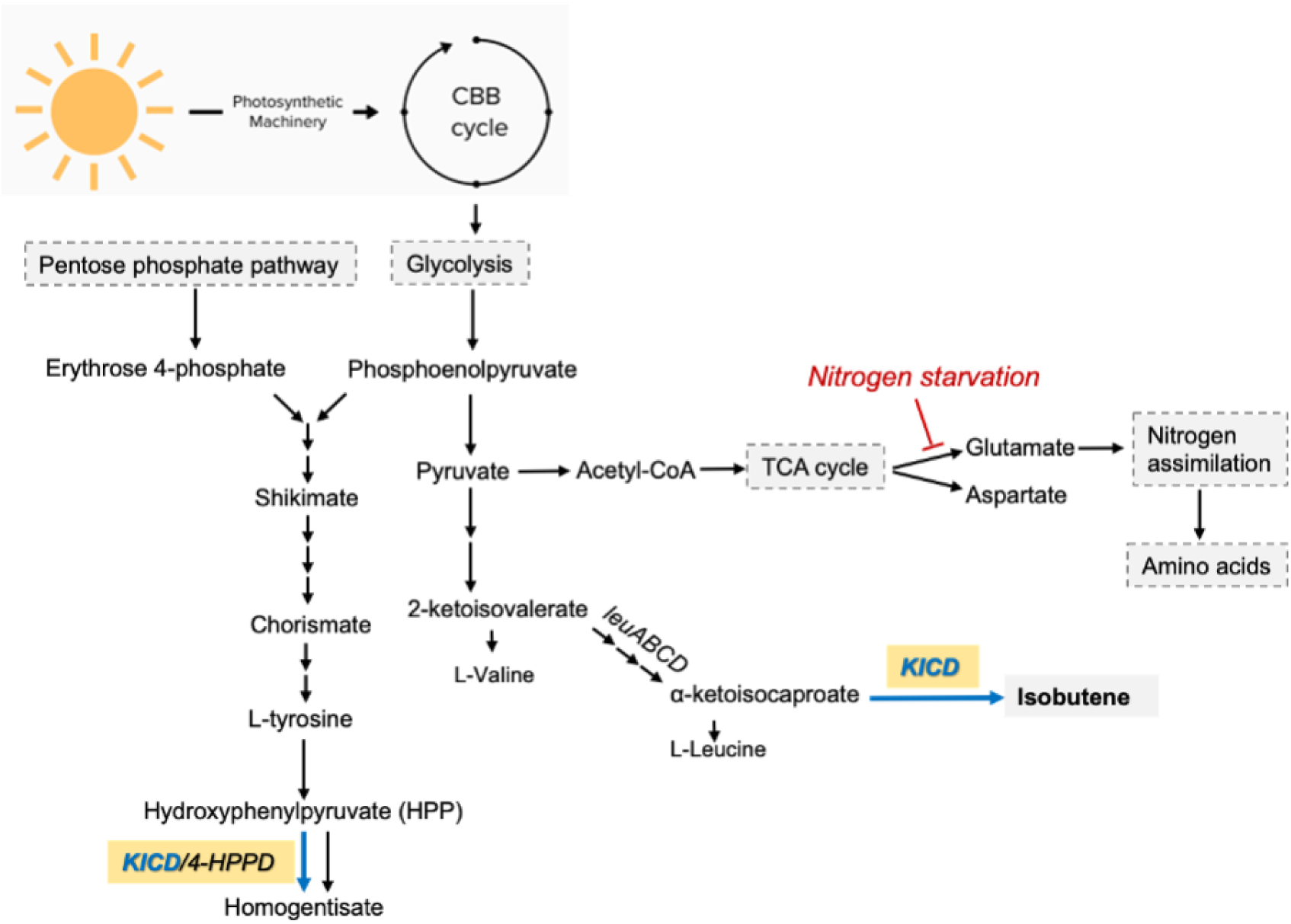
Schematic representation of *Rn*KICD’s dual role in the endogenous homogentisate and heterologous isobutene biosynthesis pathways. *Rn*KICD converts α- ketoisocaproate (KIC) into isobutene and ρ-hydroxyphenylpyruvate (HPP) into homogentisate. The black arrows indicate the native pathways, while the blue arrows represent the non-native pathways.

To further enhance isobutene production, we refreshed the cultures with BG11 medium and added 25 mM HCO₃⁻ at 408 h. Under N+ conditions, the entrapped cell beads remained stable without disintegration (Fig. 9) and continued to produce more isobutene than N-deprived cells for nearly 744 h (Fig. 10). Although the beads remained stable throughout this period, as shown by their phenotypic morphology, only a slight increase in isobutene production was observed in both entrapped Syn-F336V and Syn-*Rn*KICD cells, peaking at ∼94 mg/g DW and ∼58 mg/g DW, respectively, at 744 h (Fig. 10). These findings suggest that the six-fold increase in hydrogel-to-biomass ratio further improved bead stability and isobutene productivity by restricting cell growth, even under N+ conditions where cells tend to grow more rapidly and are more prone to cell bleaching.

To further demonstrate the effectiveness of entrapped cells’ isobutene producing capacity under N+ conditions, we compared suspension and entrapped cells with similar initial biomass loading under the same production conditions. Although up to 144 h no significant difference in isobutene productivity was observed between suspension and entrapped N+ cells (Fig. S6A), the suspension cells exhibited early bleaching as it was witnessed from 144 h onward (Fig. S6B), leading to no further increase in isobutene production. In contrast, the entrapped cells showed no signs of bleaching and continued to increase isobutene production up to 744 h, achieving productivity above 1.5- times higher than their suspension-cell counterparts (Fig. S6A). These results further confirm that cell viability and isobutene producing capacity in PVA-SA entrapped cells were enhanced as compared to the corresponding suspension cells. Further, when using entrapped cell beads to limit growth under N+ and N- conditions, we observed that N+ cells exhibited higher isobutene production than N- cells which is contrast to what we observed in the suspension cultures.

Therefore, we suggest that the nitrogen supplied entrapped cells could effectively distribute the carbon flux between BCAA biosynthesis and isobutene production, allowing for the production of essential amino acids required for cellular maintenance while achieving high isobutene output in the cells since cell growth can be limited in this case. The Syn-F336V cells showed higher isobutene production than Syn-*Rn*KICD cells (Fig.10A and B), indicating that the level of substrate available for isobutene formation is lower in the Syn-*Rn*KICD cells as compared to the Syn-F336V cells which is in line to what has been shown (Schumann et al., 2024). In contrast, under N deprived conditions, both strains experience a more pronounced metabolic shift, with carbon being redirected primarily toward isobutene production at the expense of BCAA biosynthesis. The inability to produce sufficient BCAAs under nitrogen-limited conditions impairs essential cellular functions. As a result, at ∼240 h, the strains under N- conditions experienced a sharp decline in isobutene production (Fig. 10), which could be due to reduced cell viability, correlating with their phenotypic morphology (Fig. 9), and probably due to a lack of essential amino acids. These results highlight three competing metabolic processes in the isobutene producing *Synechocystis* cells; one supporting cell growth and BCAA biosynthesis, one driving isobutene production and one producing homogentisate from HPP (Fig.11).

In the nitrogen supplied entrapped cells, both the Syn-F336V and the Syn-*Rn*KICD could maintain a balance between these three processes, resulting in prolonged cell viability and higher isobutene production, up to ∼96 mg/g DW. However, N limitation forces the cells to prioritize isobutene synthesis, at the cost of cell viability. However, this reprogramming of metabolism over time reduced productivity due to the depletion of BCAAs. This underscores the importance of nitrogen in regulating the balance between growth and isobutene production in *Synechocystis*.

## 4. Conclusions

This study demonstrated that optimizing varying environmental conditions, including light intensity and bicarbonate and nitrate concentrations, significantly enhanced isobutene production in modified *Synechocystis* cells. Under these conditions, the production rate improved over 10-fold compared to our previous report (Mustila et al., 2021). We further demonstrated that PVA-SA hydrogel-based cell entrapment was highly effective, sustaining enhanced isobutene production for nearly a month. PVA-SA hydrogel beads, crosslinked with B(OH)₄⁻ and Ca²⁺ ions, provided stability in HCO₃-rich environments, enabling long-term production. Optimization of key factors, including crosslinking time, pH balance, and the biomass-to-hydrogel ratio, improved bead durability, restricted cell growth, and prevented cell bleaching, redirecting metabolism toward isobutene production, particularly under bicarbonate and nitrate supply. Nitrogen availability emerged as a critical factor, with entrapped Syn- *Rn*KICD and Syn-F336V cell beads showing significantly higher isobutene production compared to nitrogen-limited conditions. This underscores the importance of nitrogen in balancing BCAA biosynthesis and maximizing carbon flow toward isobutene production in entrapped cells. Overall, the combination of PVA-SA hydrogel-based cell entrapment and precise nutrient management presents an effective strategy for optimizing isobutene production, providing a promising approach for sustainable bio-based chemical production.

## 5. CRediT authorship contribution statement

Sindhujaa Vajravel: Writing – original draft, Methodology, Investigation, Formal analysis, Data curation, Conceptualization. Sanjukta Aravind: Writing – original draft, Methodology, Investigation, Formal analysis, Data curation. Karin Stensjö: Writing – review & editing, Project administration, Conceptualization.

## Acknowledgements

We greatly acknowledge FORMAS—A Swedish Research Council for Sustainable Development (project no. 2021-01669), The Carl Trygger Foundation for Scientific Research (CTS:23:2849), and the Swedish Energy Agency (project no. 52576-1).

## Declaration of competing interest

The authors declare no competing financial interest.

## Appendix A. Supplementary data

Supplementary material

## Data availability

Data will be made available on request.

## Notes

### Competing Interest Statement

The authors have declared no competing interest.

### Summary of Updates

The text has been edited and improved.

## References

1. 1. Aboim, J. B., Oliveira, D. T. D., Mescouto, V. A. D., Dos Reis, A. S., da Rocha Filho, G. N., Santos, A. V., … & do Nascimento, L. A. S. (2019). Optimization of light intensity and NaNO₃ concentration in Amazon cyanobacteria cultivation to produce biodiesel. Molecules, 24(12), 2326. 10.3390/molecules24122326

2. Allen, M. M., Law, A., & Evans, E. H. (1990). Control of photosynthesis during nitrogen depletion and recovery in a non-nitrogen-fixing cyanobacterium. Archives of Microbiology, 153, 428–431. 10.1007/BF00248422

3. Allen, M. M., & Smith, A. J. (1969). Nitrogen chlorosis in blue-green algae. Archiv für Mikrobiologie, 69(2), 114–120. 10.1007/BF00409755

4. Angermayr, S. A., Rovira, A. G., & Hellingwerf, K. J. (2015). Metabolic engineering of cyanobacteria for the synthesis of commodity products. Trends in Biotechnology, 33(6), 352–361. 10.1016/j.tibtech.2015.03.009

5. Badger, M. R., & Price, G. D. (2003). CO₂ concentrating mechanisms in cyanobacteria: molecular components, their diversity and evolution. Journal of Experimental Botany, 54(383), 609–622. 10.1093/jxb/erg076

6. Bolto, B., Tran, T., Hoang, M., & Xie, Z. (2009). Crosslinked poly (vinyl alcohol) membranes. Progress in Polymer Science, 34(9), 969–981. 10.1016/j.progpolymsci.2009.05.003

7. Brouers, M., & Hall, D. O. (1986). Ammonia and hydrogen production by immobilized cyanobacteria. Journal of Biotechnology, 3(5-6), 307–321. 10.1016/0168-1656(86)90012-X

8. Carroll, A. L., Case, A. E., Zhang, A., & Atsumi, S. (2018). Metabolic engineering tools in model cyanobacteria. Metabolic Engineering, 50, 47–56. 10.1016/j.ymben.2018.03.014

9. Cheng, A. T. Y., & Rodriguez, F. (1981). Mechanical properties of borate crosslinked poly (vinyl alcohol) gels. Journal of Applied Polymer Science, 26(11), 3895–3908. 10.1002/app.1981.070261134

10. Chisti, Y. (2007). Biodiesel from microalgae. Biotechnology Advances, 25(3), 294–306. 10.1016/j.biotechadv.2007.02.001

11. Choe, S., You, S., Park, K., Kim, Y., Park, J., Cho, Y., … & Myung, J. (2024). Boric acid-crosslinked poly (vinyl alcohol): biodegradable, biocompatible, robust, and high-barrier paper coating. Green Chemistry, 26, 8230–8241. DOI: 10.1039/D4GC00618F

12. Ducat, D. C., Way, J. C., & Silver, P. A. (2011). Engineering cyanobacteria to generate high-value products. Trends in Biotechnology, 29(2), 95–103. DOI: 10.1016/j.tibtech.2010.12.003

13. Farrokh, P., Sheikhpour, M., Kasaeian, A., Asadi, H., & Bavandi, R. (2019). Cyanobacteria as an eco-friendly resource for biofuel production: a critical review. Biotechnology Progress, 35(5), e2835. 10.1002/btpr.2835

14. Forchhammer, K., & Schwarz, R. (2019). Nitrogen chlorosis in unicellular cyanobacteria–a developmental program for surviving nitrogen deprivation. Environmental Microbiology, 21(4), 1173–1184. 10.1111/1462-2920.14447

15. Forchhammer, K., & Selim, K. A. (2020). Carbon/nitrogen homeostasis control in cyanobacteria. FEMS Microbiology Reviews, 44(1), 33–53. 10.1093/femsre/fuz025

16. Gohil, J. M., Bhattacharya, A., & Ray, P. (2006). Studies on the crosslinking of poly (vinyl alcohol). Journal of Polymer Research, 13, 161–169. 10.1111/1462-2920.14447

17. Gordillo, F. J., Goutx, M., Figueroa, F. L., & Niell, F. X. (1998). Effects of light intensity, CO₂ and nitrogen supply on lipid class composition of Dunaliella viridis. Journal of Applied Phycology, 10, 135–144. 10.1023/A:1008067022973

18. Hauge, K., Bergene, E., Chen, D., Fredriksen, G. R., & Holmen, A. (2005). Oligomerization of isobutene over solid acid catalysts. Catalysis Today, 100(3-4), 463–466. 10.1016/j.cattod.2004.08.017

19. Hua, S., Ma, H., Li, X., Yang, H., & Wang, A. (2010). pH-sensitive sodium alginate/poly (vinyl alcohol) hydrogel beads prepared by combined Ca²⁺ crosslinking and freeze-thawing cycles for controlled release of diclofenac sodium. International Journal of Biological Macromolecules, 46(5), 517–523. 10.1016/j.ijbiomac.2010.03.004

20. Hulnik, M., Trofimuk, D., Nikishau, P. A., Kiliclar, H. C., Kiskan, B., & Kostjuk, S. V. (2023). Visible-light-induced cationic polymerization of isobutylene: a route toward the synthesis of end-functional polyisobutylene. ACS Macro Letters, 12(8), 1125–1131. 10.1021/acsmacrolett.3c00384

21. Kim, H. W., Vannela, R., Zhou, C., & Rittmann, B. E. (2011). Nutrient acquisition and limitation for the photoautotrophic growth of *Synechocystis* sp. PCC6803 as a renewable biomass source. Biotechnology and Bioengineering, 108(2), 277–285. 10.1002/bit.22928

22. Kosourov, S., Leino, H., Murukesan, G., Lynch, F., Sivonen, K., Tsygankov, A. A., … & Allahverdiyeva, Y. (2014). Hydrogen photoproduction by immobilized N₂-fixing cyanobacteria: understanding the role of the uptake hydrogenase in the long-term process. Applied and Environmental Microbiology, 80(18), 5807–5817. 10.1128/AEM.01776-14

23. Kostjuk, S. V. (2015). Recent progress in the Lewis acid co-initiated cationic polymerization of isobutylene and 1,3-dienes. RSC Advances, 5(17), 13125–13144. DOI: 10.1039/C4RA15313H

24. 24. Lichtenthaler, H. K. (1987). [34] Chlorophylls and carotenoids: pigments of photosynthetic biomembranes. In Methods in Enzymology (Vol. 148, pp. 350–382). Academic Press. 10.1016/0076-6879(87)48036-1

25. Lindberg, P., Park, S., & Melis, A. (2010). Engineering a platform for photosynthetic isoprene production in cyanobacteria, using *Synechocystis* as the model organism. Metabolic Engineering, 12(1), 70–79. 10.1016/j.ymben.2009.10.001

26. Markou, G., Vandamme, D., & Muylaert, K. (2014). Microalgal and cyanobacterial cultivation: the supply of nutrients. Water Research, 65, 186–202. 10.1016/j.watres.2014.07.025

27. Melis, A. (2009). Solar energy conversion efficiencies in photosynthesis: minimizing the chlorophyll antennae to maximize efficiency. Plant Science, 177(4), 272–280. 10.1016/j.plantsci.2009.06.005

28. Miller, A. G., & Colman, B. (1980). Evidence for HCO₃⁻ transport by the blue-green alga (cyanobacterium) Coccochloris peniocystis. Plant Physiology, 65(2), 397–402. 10.1104/pp.65.2.397

29. Miller, A. G., Espie, G. S., & Canvin, D. T. (1990). Physiological aspects of CO₂ and HCO₃⁻ transport by cyanobacteria: a review. Canadian Journal of Botany, 68(6), 1291–1302. 10.1139/b90-165

30. Mustila, H., Kugler, A., & Stensjö, K. (2021). Isobutene production in *Synechocystis* sp. PCC 6803 by introducing α-ketoisocaproate dioxygenase from Rattus norvegicus. Metabolic Engineering Communications, 12, e00163. 10.1016/j.mec.2021.e00163

31. Nicholas, C. P. (2017). Applications of light olefin oligomerization to the production of fuels and chemicals. Applied Catalysis A: General, 543, 82–97. 10.1016/j.apcata.2017.06.011

32. Pedruzi, G. O., Amorim, M. L., Santos, R. R., Martins, M. A., & Vaz, M. G. (2019). Biomass accumulation-influencing factors in microalgae farms. Revista Brasileira de Engenharia Agrícola e Ambiental, 24, 134–139. 10.1590/1807-1929/agriambi.v24n2p134-139

33. Peters, M. W., & Taylor, J. D. (2013). U.S. Patent No. 8,373,012. Washington, DC:U.S. Patent and Trademark Office.

34. Rana, A., Gomes, L. C., Rodrigues, J. S., Yacout, D. M., Arrou-Vignod, H., Sjölander, J., … & Ottosson, H. (2022). A combined photobiological–photochemical route to C 10 cycloalkane jet fuels from carbon dioxide via isoprene. Green Chemistry, 24(24), 9602–9619. DOI: 10.1039/D2GC03272D

35. Rebolledo-Leiva, R., Moreira, M. T., & González-García, S. (2022). Offsetting the environmental impacts of single or multi-product biorefineries from wheat straw. Bioresource Technology, 361, 127698. 10.1016/j.biortech.2022.127698

36. Rissanen, V., Vajravel, S., Kosourov, S., Arola, S., Kontturi, E., Allahverdiyeva, Y., & Tammelin, T. (2021). Nanocellulose-based mechanically stable immobilization matrix for enhanced ethylene production: a framework for photosynthetic solid-state cell factories. Green Chemistry, 23(10), 3715–3724. DOI: 10.1039/D1GC00502B

37. Rodrigues, J. S., Kovács, L., Lukeš, M., Höper, R., Steuer, R., Červený, J., … & Zavřel, T. (2023). Characterizing isoprene production in cyanobacteria–insights into the effects of light, temperature, and isoprene on *Synechocystis* sp. PCC 6803. Bioresource Technology, 380, 129068. 10.1016/j.biortech.2023.129068

38. Rossoni, L., Hall, S. J., Eastham, G., Licence, P., & Stephens, G. (2015). The putative mevalonate diphosphate decarboxylase from Picrophilus torridus is in reality a mevalonate-3-kinase with high potential for bioproduction of isobutene. Applied and Environmental Microbiology, 81(7), 2625–2634. 10.1128/AEM.04033-14

39. Sabourin, P. J., & Bieber, L. L. (1982). Purification and characterization of an alpha-ketoisocaproate oxygenase of rat liver. Journal of Biological Chemistry, 257(13), 7460–7467. DOI: 10.1016/S0021-9258(18)34400-4

40. Santos-Merino, M., Singh, A. K., & Ducat, D. C. (2019). New applications of synthetic biology tools for cyanobacterial metabolic engineering. Frontiers in Bioengineering and Biotechnology, 7, 33. 10.3389/fbioe.2019.00033

41. Sauer, J., Schreiber, U., Schmid, R., Völker, U., & Forchhammer, K. (2001). Nitrogen starvation-induced chlorosis in *Synechococcus* PCC 7942. Low-level photosynthesis as a mechanism of long-term survival. Plant Physiology, 126(1), 233–243. 10.1104/pp.126.1.233

42. Seibert, M., Kosourov, S. N., He, M., & Allahverdiyeva, Y. (2018). Immobilization of Microalgae as a Tool for Efficient Light Utilization in H2 Production and Other Biotechnology Applications (No. NREL/CH-2A00–72362). National Renewable Energy Lab. (NREL), Golden, CO (United States). 10.1039/9781849737128-00355

43. Simpson, N. E., Stabler, C. L., Simpson, C. P., Sambanis, A., & Constantinidis, I. (2004). The role of the CaCl₂–guluronic acid interaction on alginate encapsulated βTC3 cells. Biomaterials, 25(13), 2603–2610. 10.1016/j.biomaterials.2003.09.046

44. Spat, P., Klotz, A., Rexroth, S., Macek, B., & Forchhammer, K. (2018). Chlorosis as a developmental program in cyanobacteria: the proteomic fundament for survival and awakening. Molecular & Cellular Proteomics, 17(9), 1650–1669. DOI: 10.1074/mcp.RA118.000699

45. Schumann, C., Kugler, A., Berggren G., Land, H., Blikstad, C., & Stensjö, K. (2024). Structure-guided engineering of α-ketoisocaproate dioxygenase increases isobutene production in *Synechocystis* sp. PCC 6803. bioRxiv 2024-12. 10.1101/2024.12.20.629387

46. Taylor, J. D., Jenni, M. M., & Peters, M. W. (2010). Dehydration of fermented isobutanol for the production of renewable chemicals and fuels. Topics in Catalysis, 53, 1224–1230. 10.1007/s11244-010-9567-8

47. Tóth, G. S., Siitonen, V., Nikkanen, L., Sovic, L., Kallio, P., Kourist, R., … & Allahverdiyeva, Y. (2022). Photosynthetically produced sucrose by immobilized *Synechocystis* sp. PCC 6803 drives biotransformation in *E. coli*. Biotechnology for Biofuels and Bioproducts, 15(1), 146. 10.1186/s13068-022-02248-1

48. 48. Van Alphen, P., Abedini Najafabadi, H., Branco dos Santos, F., & Hellingwerf, K.J. (2018). Increasing the photoautotrophic growth rate of *Synechocystis* sp. PCC 6803 by identifying the limitations of its cultivation. Biotechnology Journal, 13(8), 1700764. 10.1002/biot.201700764

49. 49. van Leeuwen, B. N., van der Wulp, A. M., Duijnstee, I., van Maris, A. J., & Straathof, A. J. (2012). Fermentative production of isobutene. Applied Microbiology and Biotechnology, 93, 1377–1387. 10.1007/s00253-011-3853-7

50. Vajravel, S., Sirin, S., Kosourov, S., & Allahverdiyeva, Y. (2020). Towards sustainable ethylene production with cyanobacterial artificial biofilms. Green Chemistry, 22(19), 6404–6414. DOI: 10.1039/D0GC01830A

51. Valsami, E. A., Pateraki, A., Melis, A., & Ghanotakis, D. F. (2021). Heterologous β- phellandrene production by alginate immobilized *Synechocystis* sp. PCC 6803. Journal of Applied Phycology, 33, 2157-2168. 10.1007/s10811-021-02451-x

52. Wilson, J., Gering, S., Pinard, J., Lucas, R., & Briggs, B. R. (2018). Bio-production of gaseous alkenes: ethylene, isoprene, isobutene. Biotechnology for Biofuels, 11, 1–11. 10.1186/s13068-018-1230-9

53. Wu, J., Liu, R., Zhang, W., Zhong, Q., Lei, Y., & Huang, L. (2024). Enhancement of mechanical behaviors of the 3D-printed polyvinyl alcohol–based scaffold by boric acid crosslinking. Journal of Polymer Engineering, 44(4), 251–262. 10.1515/polyeng-2023-0305

54. Wu, K. Y. A., & Wisecarver, K. D. (1992). Cell immobilization using PVA crosslinked with boric acid. Biotechnology and Bioengineering, 39(4), 447–449. 10.1002/bit.260390411

55. Zhong, Y., Lin, Q., Yu, H., Shao, L., Cui, X., Pang, Q., … & Hou, R. (2024). Construction methods and biomedical applications of PVA-based hydrogels. Frontiers in Chemistry, 12, 1376799. 10.3389/fchem.2024.1376799

56. Zhu, X. G., Long, S. P., & Ort, D. R. (2008). What is the maximum efficiency with which photosynthesis can convert solar energy into biomass? Current Opinion in Biotechnology, 19(2), 153–159. 10.1016/j.copbio.2008.02.004

