## Supplementary figures for "Sustained Isobutene Production by *Synechocystis* sp. PCC 6803 Entrapped in Polyvinyl Alcohol Hydrogel Beads"

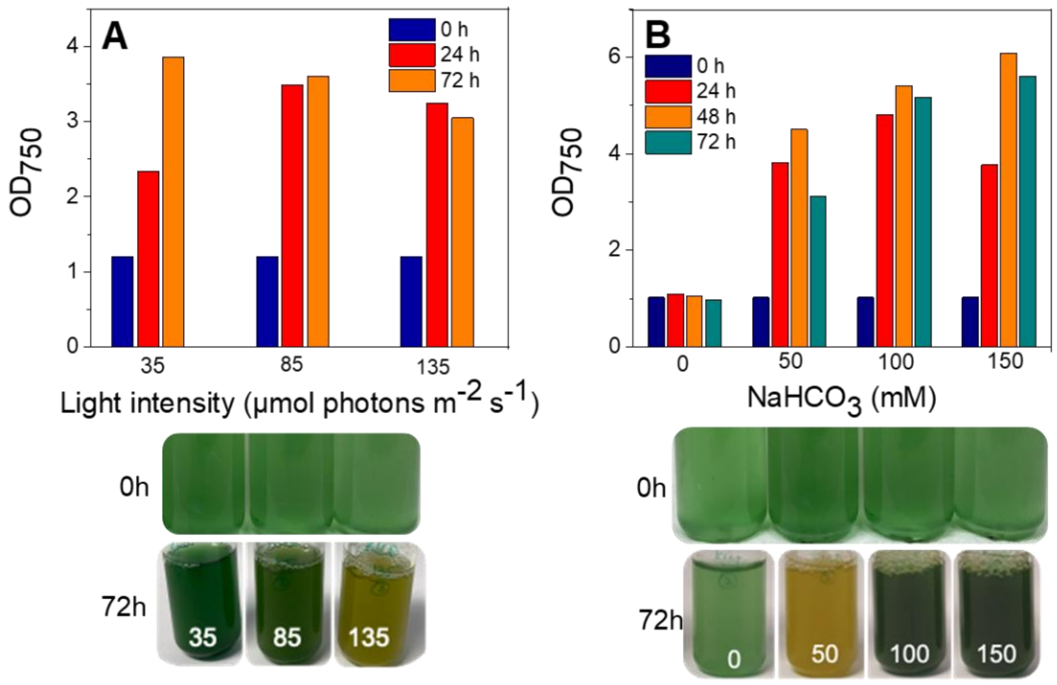


Fig S1: Cell growth and phenotypic changes of Syn-*Rn*KICD cultures under varied light intensities and inorganic carbon supplementation. A) the growth of the cells was determined by OD at 750nm alongside the phenotypic changes of cells. The cells were cultured separately under three different light intensities (35, 85, and 135 µmol photons $m^{-2}$ $s^{-1}$) while uniformly supplemented with 50 mM $NaHCO_3$. B) Influence of varying $NaHCO_3$ concentration on growth and phenotypic changes of cells under constant high light conditions (135 µmol photons $m^{-2}$ $s^{-1}$). KICD refers to Syn-*Rn*KICD.

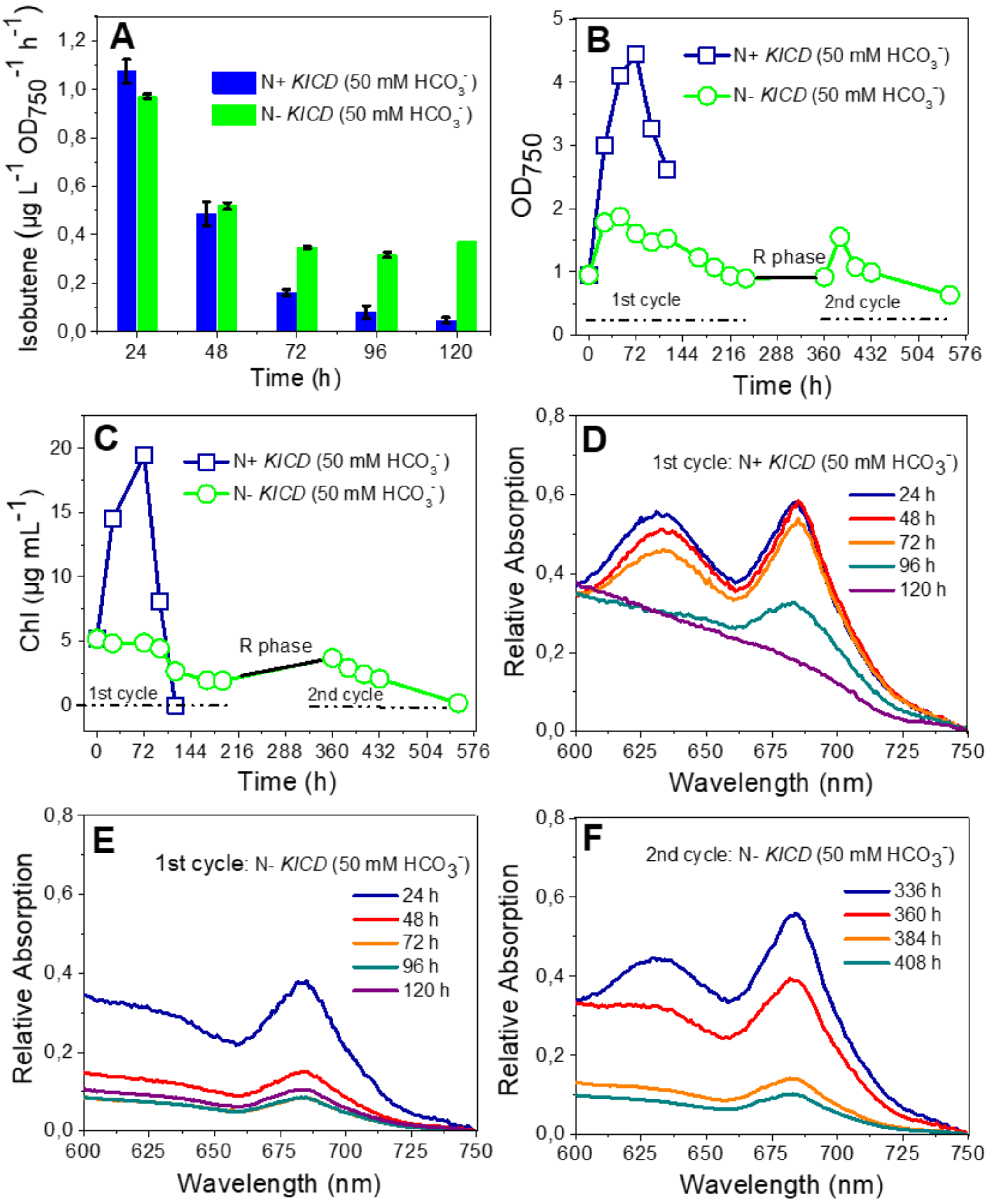


Fig S2: Effect of nitrogen starvation on isobutene productivity and major photosynthetic pigments of Syn-*Rn*KICD cells. A) The rate of isobutene productivity, B) Cell growth, and C) Chl content. D-F) ABS spectra of cells showed peaks of phycobilin ($\lambda_{max}$ = 634 nm) and Chl a ($\lambda_{max}$ = 686 nm), D) ABS spectra of N+ cells in the 1$^{st}$ cycle, E) the 1$^{st}$, and F) the 2$^{nd}$ cycle of N- cells. The whole-cell ABS spectra were measured from cells standardized to the same $OD_{750}$ values. The cultures were illuminated with 50 µmol photons $m^{-2}$ $s^{-1}$ and supplied with 50 mM $NaHCO_3$. The black arrow in Figures B and C indicates the resuscitation phase (R-Phase). KICD refers to Syn-*Rn*KICD.

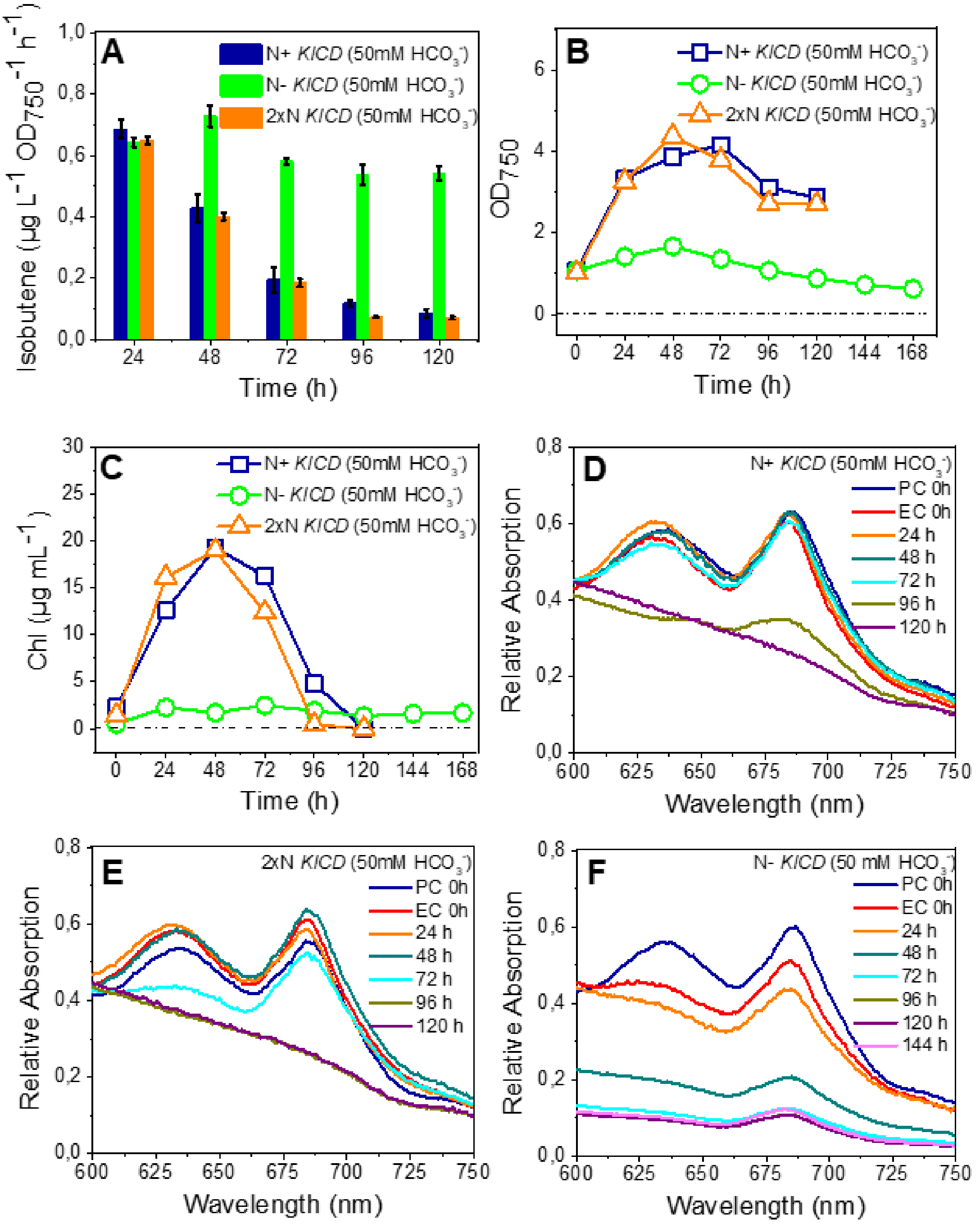


Fig S3: Effects of prolonged nitrogen starvation on the productivity of isobutene, growth and photosynthetic pigments of Syn-*Rn*KICD cells. A) The rate of isobutene productivity, B) cell grpwth, and C) Chl content. D-F) ABS spectra of cells showed PB ($\lambda_{max}$ = 634 nm) and Chl a ($\lambda_{max}$ = 686 nm), D) N+ cells, E) 2xN cells, and F) N- cells. The whole-cell ABS spectra were measured from cells standardized to the same $OD_{750}$ values. The cultures were illuminated with 50 µmol photons $m^{-2}$ $s^{-1}$ and supplied with 50 mM $NaHCO_3$. In Figure D-F, PC refers to the cultures prior to the N starvation, and EC refers to the experimental cultures after N starvation. KICD refers to Syn-*Rn*KICD.

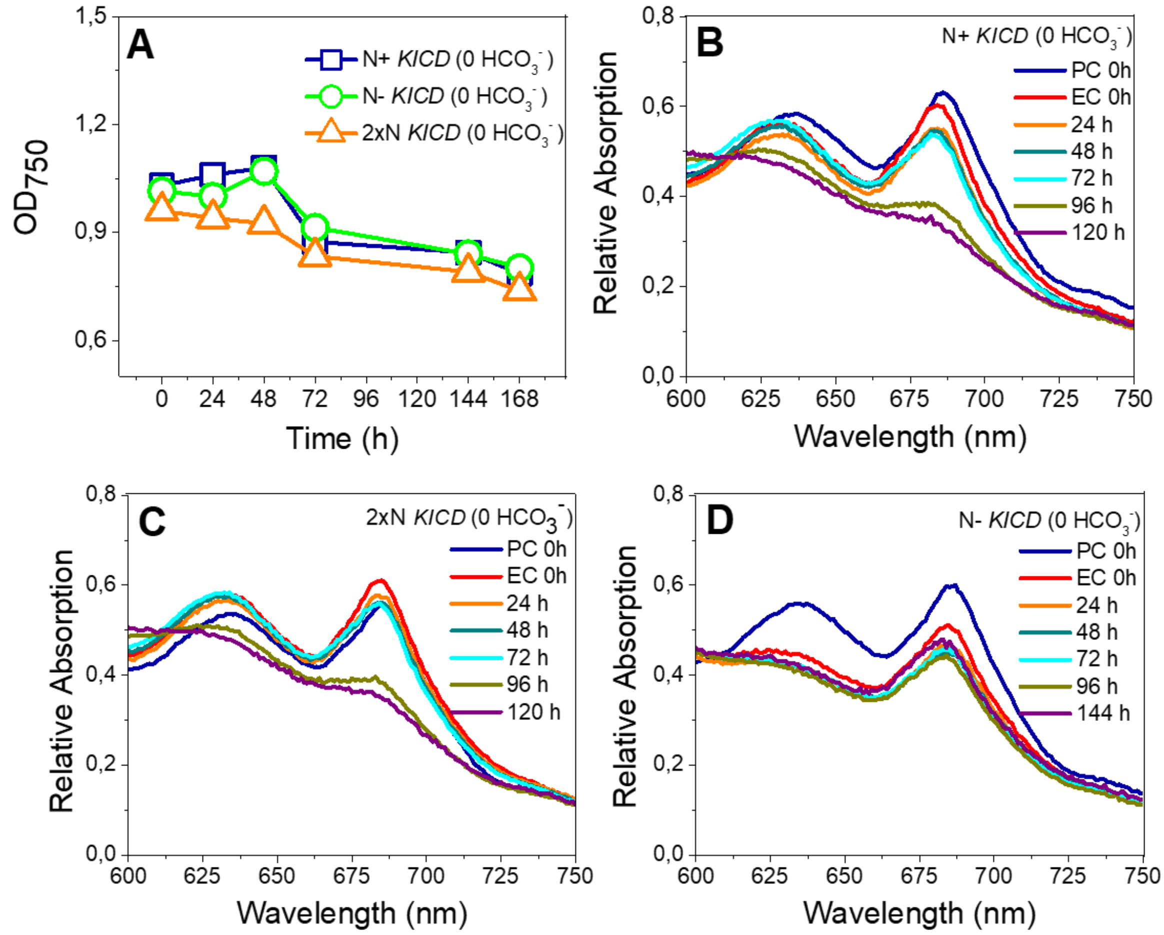


**S4:** Effects of both nitrogen and carbon limitation on Syn-*Rn*KICD cell growth and photosynthetic pigments. A) cell growth (OD750), B-D) ABS spectra of cells under N+, 2xN, and N- conditions. All the cultures were deprived of $HCO_3$ and illuminated with 50 μmol photons $m^{-2}$ $s^{-1}$. In Figure B-D, PC refers to the cultures prior to the N starvation, and EC refers to the experimental cultures after N starvation. KICD refers to Syn-*Rn*KICD.

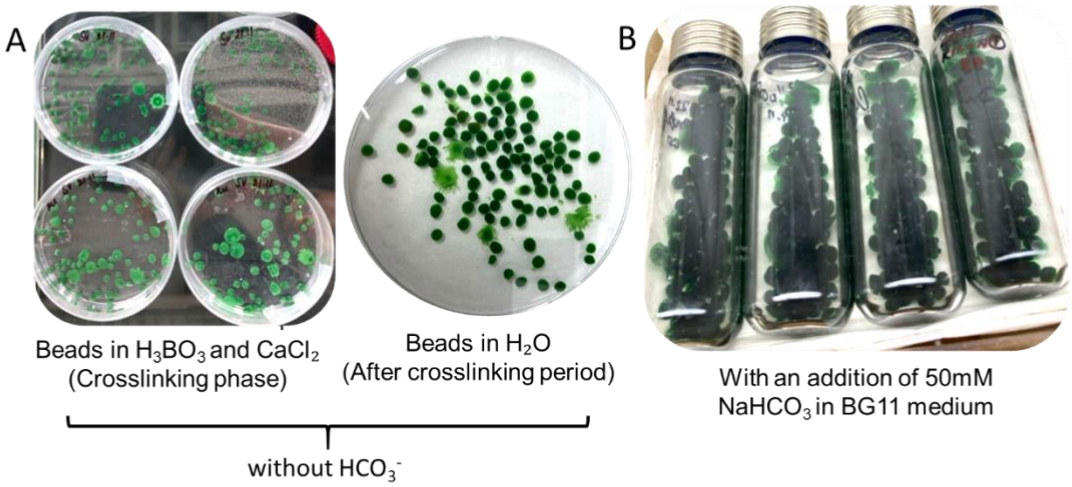


**S5:** Enhanced bead stability in Syn-*Rn*KICD cells with a 2-fold higher PVA-SA hydrogel-entrapped cell system**.** A) Physical characteristics of beads during crosslinking phase without the addition of $NaHCO_3$ under 30 µmol photons $m^{-2}$ $s^{-1}$. B) After the crosslinking period, the cells were supplemented with 50 mM $NaHCO_3$ in tightly sealed vials and illuminated with 50 µmol photons $m^{-2}$ $s^{-1}$. The beads were prepared using a 5:12:1 ratio of cell mass to hydrogel (PVA-SA) and subjected to a 20-hour crosslinking process with 4% $H_3BO_3$ and 2% $CaCl_2$. KICD refers to Syn-*Rn*KICD.

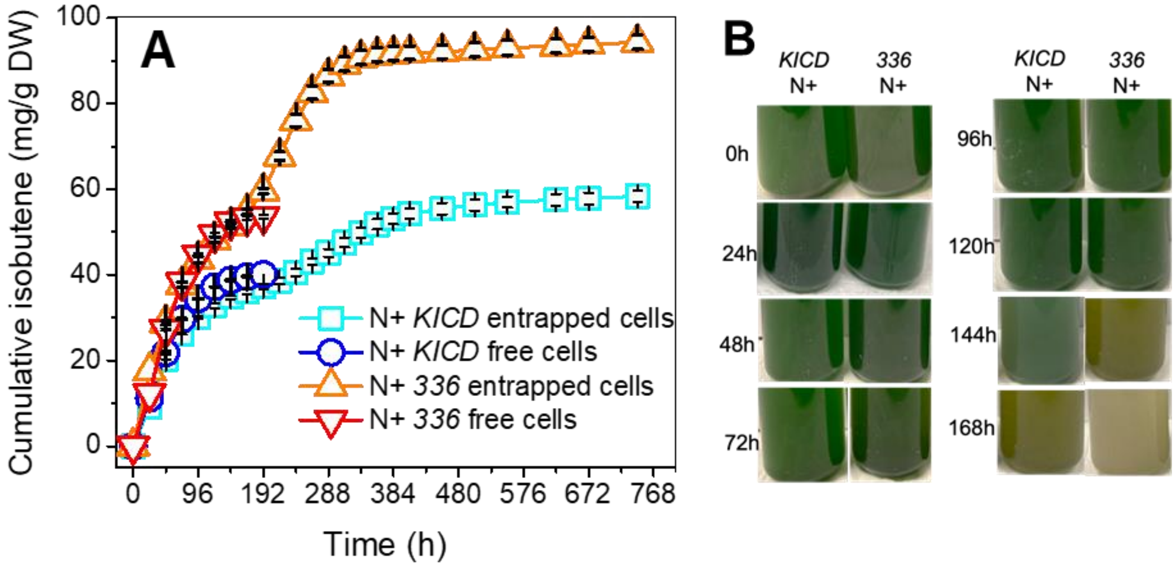


**S6:** Comparison of isobutene production in cell suspension and entrapped Syn-*Rn*KICD and Syn-F336V cells under different nitrogen supplementation regimes**.** A) the cumulative isobutene productivity and B) the phenotypic changes of cells. A similar amount of initial biomass was used to compare the isobutene productivity between suspension and immobilized cells. pH was adjusted to neutral every 24 h in all the culture conditions, including suspension and entrapped cells. The beads were prepared using a 5:36:3 ratio of cell mass to hydrogel (PVA-SA). The entrapped cells were supplemented with 50 mM $NaHCO_3$ and illuminated with 50 μmol photons $m^{-2}$ $s^{-1}$. The productivity of isobutene was normalized to the cells' initial DW. In the figure, 336 refers to Syn-F336V, and KICD to Syn-*Rn*KICD
